# Post-stroke fatigue as a disorder of uncoupled agency: an exploration of predictive and postdictive mechanisms

**DOI:** 10.64898/2026.07.08.737231

**Authors:** Massimo Bertoli, William De Doncker, Annapoorna Kuppuswamy

## Abstract

Post-stroke fatigue is a common and disabling consequence of stroke, defined by a chronically elevated sense of effort during voluntary action that is disproportionate to any objective motor impairment. The sensory attenuation framework proposes that this effort distortion reflects reduced but not abolished gain on descending motor predictions. Because the same gain is thought to underlie agency, the framework predicts that in post-stroke fatigue peripheral sensory attenuation should be preserved, the cortical predictive signal should be disrupted, and postdictive integration should be amplified. We tested these predictions using two implicit measures of agency probing distinct levels of the sensorimotor hierarchy.

Data were drawn from two independent cross-sectional studies of stroke survivors (≥3 months post-stroke), classified as high (Fatigue Severity Scale-7 ≥ 5) or low fatigue. A force matching task, indexing peripheral forward-model attenuation, was analysed in 40 patients (30 low-fatigue, 10 high-fatigue) and 24 healthy controls. An intentional binding paradigm with concurrent EEG, indexing the cortical predictive (readiness potential) and postdictive (outcome binding) components of agency, was completed by 23 patients (15 low-fatigue, 8 high-fatigue), of whom 17 yielded analysable EEG. Mixed-effects models were the primary framework, with fatigue group as the principal predictor, anxiety and depression as covariates, and robustness assessed via permutation, leave-one-out, Bayesian and equivalence analyses.

Sensory attenuation was robust (n = 64) and unmodulated by fatigue, with the smallest detectable interaction small relative to mean attenuation. Outcome binding was selectively amplified in high-fatigue patients, showing temporal compression 2.5-fold greater than low-fatigue patients (72.0 vs 28.6 ms), whereas action binding was unmodulated (three-way interaction P = 0.007). The readiness potential showed canonical agency-related enhancement in low-fatigue but not high-fatigue patients (interaction P = 0.0002, robust to permutation, leave-one-out and Bayesian analyses), though the high-fatigue finding (n = 5) is preliminary.

A direct test of predictive-postdictive uncoupling in the 17 patients with both measures was non-significant; uncoupling is therefore advanced as a working hypothesis, not an established finding. All effects were robust to mood adjustment.

These findings are consistent with preserved peripheral attenuation, a disrupted cortical predictive signal, and amplified postdictive integration in post-stroke fatigue, compatible with a single upstream reduction in descending predictive gain and, tentatively, a shift from diminished to uncoupled agency. This differs from the peripheral-attenuation failure of schizophrenia and functional movement disorders, suggesting an intermediate position within the agency-disorder spectrum, and points to the cortical predictive layer as a candidate target for future work.

## Introduction

The sense of agency refers to the experience of being the author of one’s own actions and the events they produce, and represents a basic and constant feature of voluntary motor behaviour^1^. Although a unitary experience, literature has established two dimensions that contribute to the sense of agency. The first is a phenomenological distinction: on the one hand, a non-conceptual *feeling of agency* referring to the pre-reflective sense that an ongoing action is one’s own. On the other hand, a conceptual *judgement of agency*, that is, an a posteriori attribution of authorship. The two levels are dissociable: one may experience an intact feeling of agency while making delusional judgements about authorship, or, conversely, retain accurate judgement while showing altered implicit attribution. The second dimension is a temporal-source distinction: it has been advanced that authorship develops through the optimal integration of cues coming from both predictive signals available before and during the action (e.g., efference copies of motor commands) and postdictive signals available after the action (e.g., sensory feedback, affective valence of the outcomes)^2^. Crucially, like the phenomenological levels above, these two sources are dissociable as well: pathology can selectively degrade the predictive contribution while sparing or even forcing greater reliance on the postdictive one, shifting the balance of cue integration rather than abolishing agency ^2,3^.

Two implicit indices have been established as the chief experimental tools for probing the feeling of agency level. First, sensory attenuation (SA) refers to the systematic reduction in perceived intensity of self-generated reafference compared to externally produced inputs of equivalent magnitude ^4,5^, although the perceptual interpretation of such attenuation effects remains debated ^6^. Because the motor system uses an internal forward model to predict the sensory consequences of its own commands ^7^ and attenuates the input that matches this prediction, SA indexes predominantly the predictive source of agency ^8^. Second, intentional binding (IB) refers to the implicit temporal compression of the perceived time intervals between voluntary actions and their sensory consequences ^9,10^. The effect is measured using the Libet clock paradigm, i.e., a rotating-clock display originally developed by Libet et al. ^11^ to capture the subjective timing of voluntary intention, and subsequently adapted by Haggard et al. ^9^ to assess the perceived timing of self-generated actions and their sensory outcomes. Moreover, IB provides dissociable action-binding and outcome-binding components that index, respectively, predictive and postdictive sources ^2,12^ with action binding depending on efferent, prediction-based signals ^13^. SA and IB are therefore complementary windows onto the predictive-postdictive cue integration that underlies the feeling of agency, accessed without the response bias that affects explicit measures of agency judgement ^9^. At the computational level, this cue integration can be expressed as a precision-weighted comparison of descending motor predictions with afferent sensory feedback. Under active-inference accounts of motor control ^14,15^, the motor system generates predictions about the sensory consequences of its commands and applies a high-precision gain function to these descending signals, attenuating predicted reafference and labelling it as self-generated. SA is the direct perceptual marker of this attenuation, while IB is its temporal-perceptual counterpart, arising when the action is integrated with its effect and drawing on both predictive and inferential contributions ^16^. The same gain operation has a third, complementary output. Where SA reflects the reafference that is successfully attenuated, the sense of effort reflects what is left unattenuated. In keeping with this active-inference account, perceived effort during voluntary action is not a direct read-out of peripheral afferent feedback from the working muscles ^17,18^ but tracks the prediction error remaining after descending attenuation ^14,19^. Precise predictions produce small residual errors and low perceived effort, while imprecise predictions produce large residual errors and elevated perceived effort. Sensory attenuation, intentional binding, and effort can be seen, therefore, not as independent phenomena but as three perceptual read-outs of a single underlying computation: the precision-weighted matching of predicted and inferred actual reafference. A consequence follows directly. Because these read-outs share one generative mechanism, they should not change independently, and a disturbance of that mechanism is expected to produce a coordinated but uneven pattern across them. In particular, a partial reduction in central gain can spare the peripheral read-out while reshaping the central and postdictive ones, resulting in a distinctive profile that reveals where the deficit lies.

Empirical illustration of this coupling comes from a family of clinical conditions collectively termed “*disorders of agency*” in which the integrity of descending predictive gain is compromised to varying degrees. At one end of this spectrum, schizophrenia provides the clearest existing demonstration of severe SA failure: in the force matching task (FMT), in which healthy participants reproduce an externally applied force and self-generated forces are systematically overestimated, patients instead show reduced overestimation indexing impaired peripheral SA at the behavioural level ^20^, together with altered processing of effort during voluntary action ^21,22^ and the categorical experience that one’s own actions are caused by external agents ^23^. Functional movement disorders show a parallel pattern: reduced SA in the force matching task ^24^ coupled with disrupted agency over symptomatic movements. Both scenarios are interpreted as a near-abolition of descending predictive gain, that is, predictions too imprecise to attenuate reafference, which is then processed as exafference (i.e., as if externally generated) and ultimately misattributed ^14,25^. Within the cue-integration framework ^2^, the loss of precise predictive cues forces a disproportionate reliance on postdictive ones, which in schizophrenia can produce delusional attribution. At an intermediate point along this same spectrum, theoretical accounts predict a distinct phenotype in which descending predictive gain is degraded but not abolished. Its predicted phenomenological signature differs qualitatively: heightened rather than misattributed effort, and distorted rather than absent agency, all within preserved categorical self-attribution ^19^. To our knowledge, the agency-computational predictions entailed by this intermediate phenotype have never been directly tested in any clinical population.

Post-stroke fatigue (PSF) stands as the most prevalent candidate for this intermediate phenotype. Affecting approximately half of stroke survivors and persisting for years after the acute event, chronic PSF is defined phenomenologically by a chronically elevated sense of effort during voluntary action that is disproportionate to any objective motor impairment, in line with the classical definition of central fatigue in neurological conditions ^26^, with task performance and reported fatigue levels characteristically uncoupled ^19,27^. Lesion location is not considered a primary determinant of chronic fatigue ^27^. The sensory attenuation framework (SAF) of pathological fatigue ^19,28^ proposes that this effort distortion may reflect reduced but not abolished gain on descending motor predictions, eventually yielding abnormally inflated residual prediction errors. Direct experimental support has linked the subjective experience of fatigue to the motor system at progressively more specific levels. At the perceptual level, higher Fatigue Severity Scale scores (FSS-7; ^29–31^) are associated with elevated implicit perceived effort, indexed by a bias on a concurrent line-length judgement task ^32^. At the level of corticomotor physiology, high fatigue is accompanied by reduced excitability of the corticospinal output and of its facilitatory inputs ^33^. This hypo-excitability has a candidate network origin in the shifted interhemispheric inhibitory balance between homologous primary motor regions that predicts fatigue severity ^34^, and it becomes manifest, trial by trial, as a failure to reach the appropriate pre-movement state during movement preparation ^35^. Across these levels, however, the deficit has been characterised exclusively in terms of motor excitability and preparatory state. Whether the same impairment extends to the computation of agency itself, or in other terms whether the brain’s capacity to predict the sensory consequences of voluntary action, and to bind those consequences to the action that caused them, is altered in PSF, has never been tested.

The same precision-weighted comparison is instantiated at successive levels of this hierarchy, each accessible through a distinct measure: a peripheral, effector-level forward model indexed by FMT overestimation; a cortical, preparatory predictive stage indexed by the Readiness Potential (RP); and a postdictive integration stage indexed by outcome binding. Descending predictive gain that is reduced but not abolished should therefore leave a level-specific rather than uniform signature: preservation at the peripheral level, disruption of the cortical predictive stage, and compensatory amplification of postdictive integration. From this, several predictions follow. At the peripheral level, effector-level SA should be preserved. In PSF, the SAF-posited reduction in predictive gain has been localised to central levels of the motor hierarchy ^33–35^ and is computationally dissociable from the peripheral effector forward model indexed by the FMT. A partial, central reduction can drive effort and disrupt cortical preparation while sparing peripheral attenuation, so that preserved peripheral SA is the specific prediction for an intermediate phenotype. In contrast, a peripheral failure would align PSF with the abolished-gain end of the spectrum (schizophrenia, functional movement disorder). Moreover, the postdictive component (the retrospective integration of action consequences into the experience of agency), should be amplified, because under the cue-integration scheme the system compensates for degraded predictive cues by upweighting postdictive ones; this is further grounded by Demanet et al. ^36^, who showed that task-unrelated physical effort amplifies IB in healthy participants, demonstrating that effort signals enter the postdictive cue pool. Finally, the predictive component (the pre-motor preparatory activity that anticipates the sensory consequences of voluntary action), should be selectively disrupted in high-fatigue patients. The electrophysiological correlate of this activity is the RP, first described by Kornhuber & Deecke ^37^ and generated in the pre-supplementary motor area (pre-SMA) ^38^. Although historically conceptualised as a generic motor-preparation signal, the RP has been experimentally re-established as a correlate of agency-related computation rather than of motor execution per se: transcranial magnetic stimulation of the pre-SMA, a principal cortical generator of the RP, selectively reduces IB, whereas theta-burst stimulation of the sensorimotor cortex does not ^39^, and RP amplitude correlates within subjects with the magnitude of IB ^40^. The RP therefore provides direct neurophysiological access to the descending predictive signal that the SAF identifies as compromised in PSF.

To test these predictions, we drew on two independently conducted studies on chronic stroke survivors, of complementary anatomical and computational scope: a FMT, which indexes peripheral, effector-level forward-model attenuation, and an IB paradigm with concurrent EEG recording of the RP, which indexes the cortical-predictive and postdictive components of the same hierarchy. The two paradigms are validated measures of voluntary agency: the FMT through the contrast between self-generated and externally produced forces of identical magnitude ^5^, the IB through the reversal of temporal compression when the same movement is produced involuntarily by transcranial magnetic stimulation ^9^. Examined together across comparable patient cohorts, they allow the predictive and postdictive components of agency, and the peripheral and cortical levels of the predictive hierarchy, to be dissociated within a single theoretical framework. While IB has been applied to schizophrenia ^3^, functional movement disorders ^41^, and Tourette syndrome ^42^, to our knowledge no published study has applied it to PSF or to any chronic fatigue condition.

In two cohorts of chronic stroke survivors classified as high (FSS-7 ≥ 5) or low (< 5) fatigue ^33^, we accordingly test three predictions of the SAF account of PSF as an agency disorder: (i) that peripheral SA, indexed by FMT overestimation, is preserved across the FSS range, thus distinguishing PSF mechanistically from the peripheral SA failure phenotype of schizophrenia and functional movement disorders; (ii) that outcome binding is amplified in high-fatigue patients; (iii) that the early RP fails to differentiate operant from baseline conditions in high-fatigue patients. Jointly, this dissociation would constitute the first integrated empirical test of the SAF account across multiple levels of the agency hierarchy and would establish PSF as an intermediate phenotype within the broader spectrum of agency-related disorders.

## Materials and methods

### Participants

The present manuscript combines data from two independent cross-sectional observational studies. The two studies were not designed to be performed in sequence: the force matching task (FMT) study and the intentional binding (IB) study employed different cohorts assessed in separate sessions, with a convenience overlap of 14 patients participating in both. Both studies were approved by the London Bromley Research Ethics Committee (REC reference number: 16/LO/0714). Following written informed consent in accordance with the Declaration of Helsinki. Two cohorts of chronic stroke survivors were recruited and contributed to this study: 23 chronic stroke survivors who completed the IB task with EEG (henceforth the IB cohort) and 46 chronic stroke survivors recruited for the FMT (FMT cohort). In addition, 24 healthy controls (HC) with no history of neurological disease completed the FMT and served as a normative reference for the FMT analysis. After quality control (Supplementary Methods 2), the primary FMT analysis included 40 patients (30 low-fatigue, 10 high-fatigue) and the 24 HC. The IB analysis included all 23 patients. For both IB and FMT cohorts, stroke survivors were recruited via the Clinical Research Network from the University College Hospital NHS Trust (UCLH), the departmental Stroke Database and the community. Eligibility criteria were: (i) first-ever ischaemic or haemorrhagic stroke; (ii) at least 3 months post-stroke at testing; (iii) no other clinical neurological diagnosis; (iv) absence of moderate-to-severe depression (Hospital Anxiety and Depression Scale Depression subscale ≤ 11; ^43^) and not taking antidepressants or other centrally acting medication; (v) physical recovery sufficient to perform the task, operationalised as grip strength and Nine-Hole Peg Test performance of the affected hand ≥ 60 % of the unaffected hand. Demographic characteristics of stroke survivors cohorts are summarised in Table 1.

**Table 1.** Demographic and clinical characteristics of the Intentional Binding (IB) and Force Matching Task (FMT) cohorts, stratified by fatigue group.

| Variable | IB cohort (n = 23) |  |  | FMT cohort (n = 46) |  |  | Between cohorts *P* |
| --- | --- | --- | --- | --- | --- | --- | --- |
|  | LowFSS<br>(n = 15) | HighFSS<br>(n = 8) | *P* | LowFSS<br>(n = 31) | HighFSS<br>(n = 15) | *P* |  |
| Age (years) | 65.3<br>(55.3–69.9) | 60.6<br>(52.1–69.3) | 0.821 | 64.0<br>(58.0–71.0) | 56.0<br>(48.0–67.0) | 0.128 | 0.934 |
| FSS-7 | 3.3<br>(1.8–3.6) | 5.8<br>(5.4–6.3) | <0.001 | 2.4<br>(1.6–3.6) | 5.7<br>(5.1–6.0) | <0.001 | 0.554 |
| HADS-Depression | 5.0<br>(2.0–8.0) | 7.5<br>(3.0–9.0) | 0.381 | 3.0<br>(2.0–5.5) | 7.0<br>(5.0–9.0) | 0.005 | 0.518 |
| HADS-Anxiety | 3.0<br>(2.0–5.5) | 7.0<br>(2.8–10.0) | 0.193 | 4.0<br>(2.0–6.0) | 6.0<br>(4.0–8.0) | 0.141 | 0.458 |
| Time post-stroke (y) | 7.2<br>(5.2–9.5) | 9.1<br>(6.7–16.4) | 0.165 | — | — | — | — |
| Hemisphere (L / R) | 8 / 7 | 4 / 4 | 1.000 | 14 / 17 | 7 / 8 | 1.000 | 0.621 |

Values are median (interquartile range) unless otherwise indicated. LowFSS, Fatigue Severity Scale (7-item) score < 5; HighFSS, FSS-7 ≥ 5. Within-cohort comparisons (LowFSS vs HighFSS) and between-cohort comparisons (same variable across IB and FMT) used the Mann–Whitney U test for continuous variables and Fisher's exact test for hemisphere. FSS-7, Fatigue Severity Scale, 7-item version; HADS-D, Hospital Anxiety and Depression Scale, Depression subscale; HADS-A, HADS, Anxiety subscale.

### Fatigue assessment

Trait fatigue, defined as the experience and functional impact of fatigue in the two weeks preceding testing, was quantified using the 7-item Fatigue Severity Scale (FSS-7; ^31^). State fatigue, the momentary subjective sense of tiredness on the day of testing, was measured using a visual analogue scale ranging from 0 (“not at all tired”) to 10 (“extremely tired”) in unit-point steps. Patients were classified into LowFSS (FSS-7 < 5) and HighFSS (FSS-7 ≥ 5) subgroups^44^. Hospital Anxiety and Depression Scale anxiety and depression subscale scores (HADS-A and HADS-D) were collected as covariates for sensitivity analyses.

### Force Matching Task

#### Apparatus and stimuli

Force matching was performed on a purpose-built device developed for use in stroke survivors. Standard robotic rigs require the stimulated hand to be held supine throughout testing, a posture that may become uncomfortable over a prolonged session. In the present device, the hand receiving the force instead rested on a cushion in a neutral posture, midway between pronation and supination. The same device served as both force generator and force sensor. Force was produced pneumatically: regulated airflow delivered under pulse-code modulation set the magnitude transmitted to the finger, with air pressure determining the force level (Fig. 1, top). Command levels were driven through the digital and analogue channels of a Power1401 acquisition unit (CED) running Signal (v6.04, CED), and the force-sensor signal was acquired at 2,500 Hz through the unit’s digital inputs and monitored in Signal. The system was calibrated such that a 0.25 V command output corresponded to 1 N of applied force, registering as a 0.0197 V reading at the force sensor. Sensor readings were therefore converted to newtons (N) by dividing by this factor.

**Figure 1.**
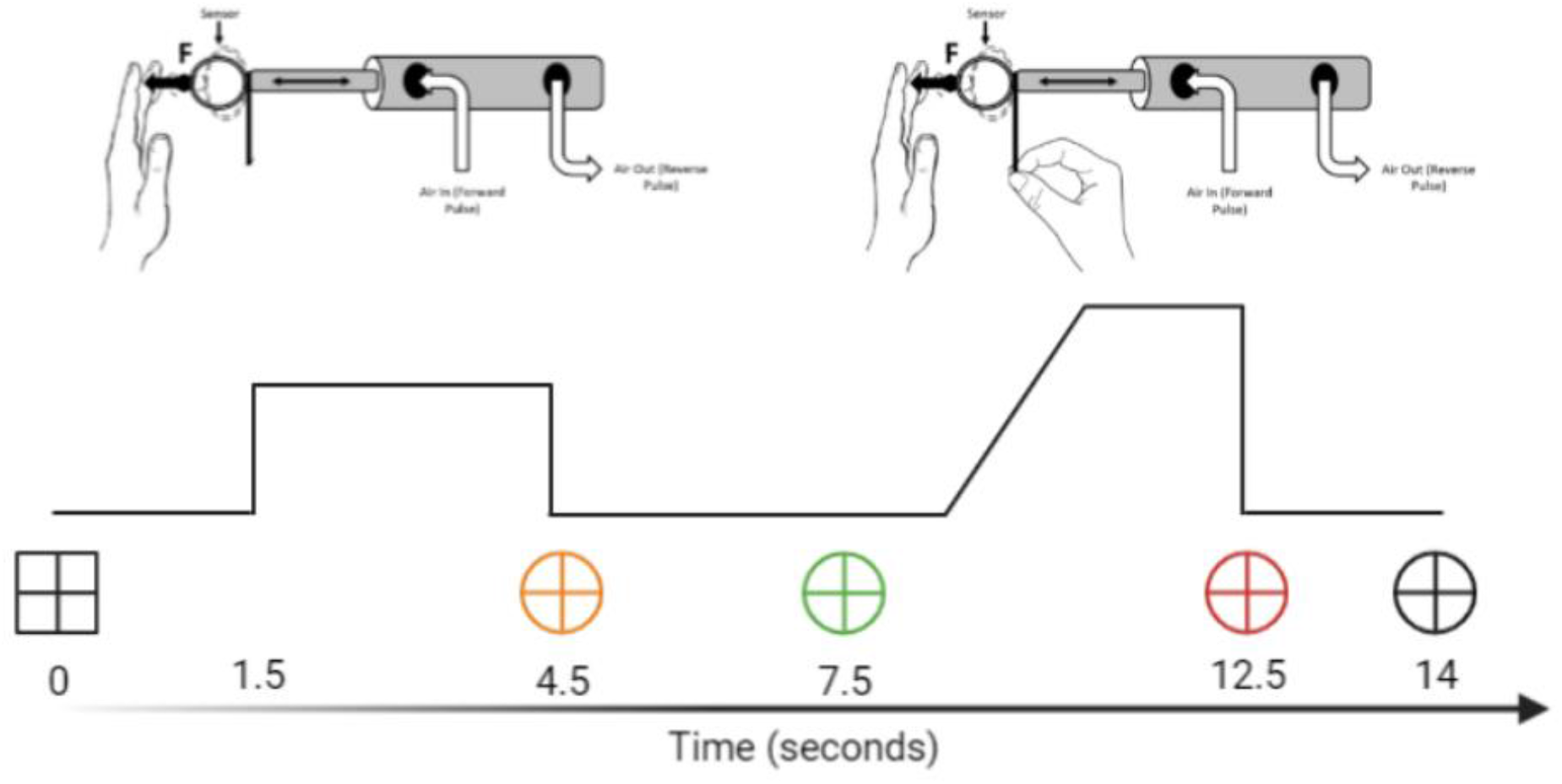
Force matching task apparatus and trial structure. Participants matched a target force delivered to the left index finger by a torque motor (the *applied* force) by pressing on a force transducer with the right index finger (the *matched* force), without visual feedback. The schematic (top) shows the apparatus, with the lever and force sensor through which the applied force is generated and the matched force recorded. The time course of a trial (bottom) illustrates the sequence of target force levels, with the applied force held at each of the target levels (1, 2 and 3 N; the 20 N level was excluded from the primary analysis) over the trial period (seconds)

#### Procedure

Participants sat at a table with the tip of the left index finger on a moulded rest at the transducer, facing a computer monitor (DELL 1909W, 19” LCD Display). Each trial started with a black box and central fixation cross. 1.5 seconds later, the transducer applied the target force to the left index finger for 3 seconds. Target forces were drawn from four levels (1, 2, 3, and 20 N) in pseudo-random order, each occurring once per four-trial cycle. The force was then withdrawn, and an orange circle with a fixation cross signalled a 3-second retention interval, during which participants held the just-experienced force in memory without responding. A green circle then cued the matching phase: using the right hand, participants reproduced the experienced force by manipulating a lever on the transducer, with up to 5 seconds to reach, and were instructed to hold that force level until a red’stop’ circle appeared, whereupon they released the lever and the transducer reset automatically. A black circle marked the 1.5-second inter-trial interval. Fig. 1 (bottom) depicts a visual representation of the time course and structure of a single trial. The sensor recorded both applied (target) and matched forces at the left index finger. Participants first completed a 12-trial familiarisation phase (three cycles of the four forces). The main task comprised eight blocks of 20 trials (160 trials), with five trials per force level per block so that total force was equated across blocks, and a two-minute rest between blocks. No applied force was reported as painful. Stimuli were delivered by a bespoke Signal (v6.04, CED) script.

#### Outcome measures

Sensory attenuation (SA) was operationalised as the systematic overestimation of matched relative to applied force (F) as follows:

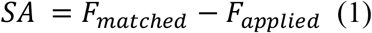

This index reflects the failure of the forward-model attenuation mechanism that, in healthy individuals, calibrates the perceived intensity of self-generated reafference ^4,5^. A linear fit was computed per participant relating matched to applied force across States 1, 2 and 3 N, yielding two secondary indices: the *gradient* (proportional scaling of overestimation with target force) and the *intercept* (baseline overestimation at zero force). Trials at the 20 N force level were excluded from primary analyses owing to high inter-trial and inter-subject variability arising from apparatus saturation at this force level, and the validity of this exclusion was verified in a sensitivity analysis (see Results). Subject-level data quality was assessed by trial completeness per force level: blocks with fewer than 50 % of expected trials at any of the analysed force levels (1, 2 or 3 N) were excluded. A more conservative criterion (≥ 75 % trial completion plus exclusion of blocks with coefficient of variation in matched force exceeding the cohort mean + 3 SD) was applied in a sensitivity analysis (Supplementary Methods 2, Supplementary Table S1).

### Intentional Binding Task

#### Apparatus and stimuli

Participants were seated 70 cm from a 24-inch monitor (Dell U240, 1280 × 768 resolution) and responded via USB keyboard. The experiment was controlled by the Psychophysics Toolbox^45^ running on a Windows computer. Auditory stimuli were delivered through headphones (Sennheiser HD 569).

The timing of perceived events was measured using the adapted Libet clock paradigm ^9,11^ for the assessment of IB. The clock display consisted of a circular dial with 60 tick marks numbered at 5-unit intervals (5, 10, 15,…, 60). A single clock hand of unpredictable starting position completed a full rotation every 2560 ms, yielding a temporal resolution of approximately 42.7 ms per clock unit. On each trial, participants verbally reported the perceived clock-hand position as an integer from 1 to 60, which the experimenter recorded (Fig. 2, Panel B).

**Figure 2.**
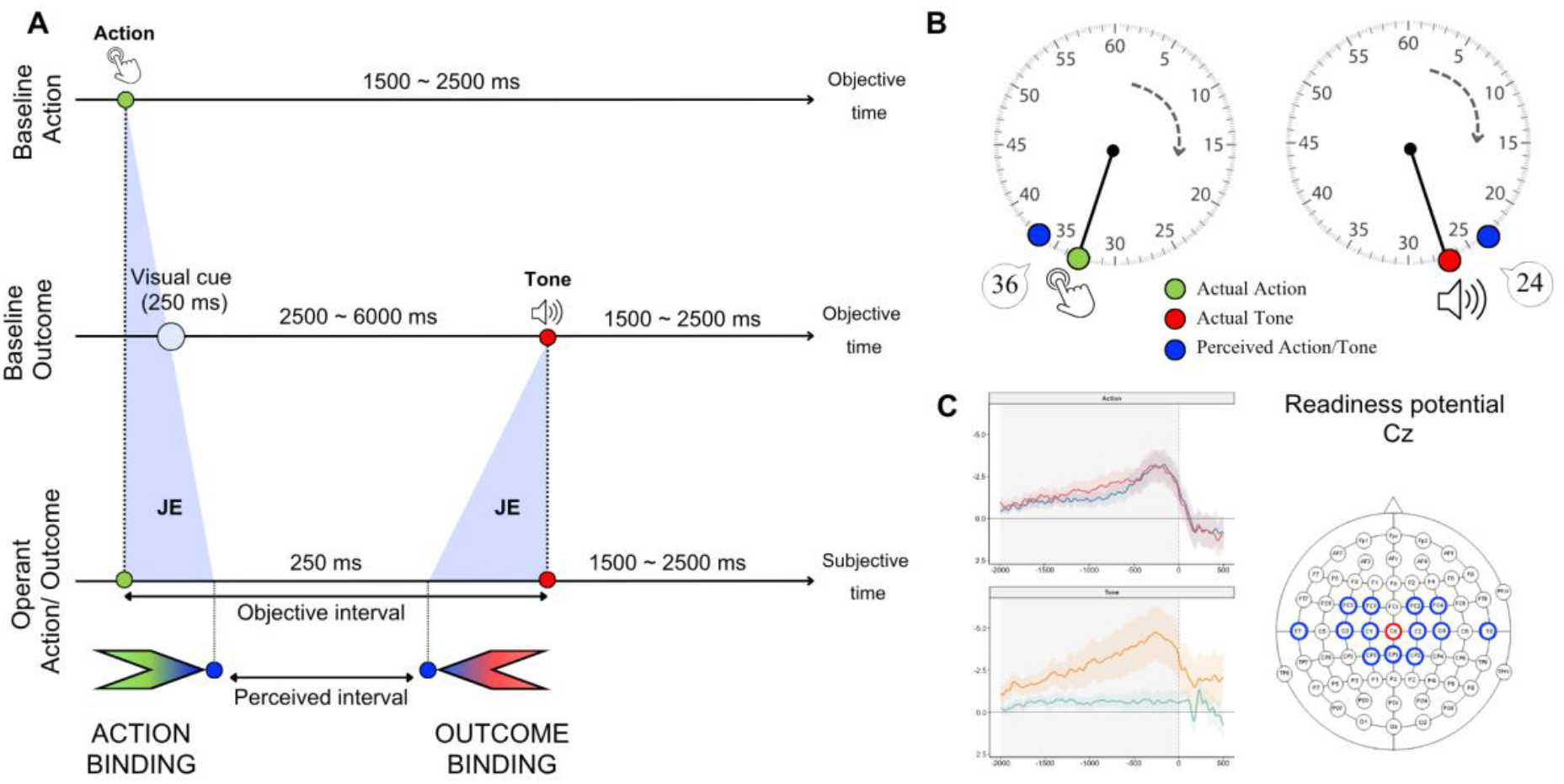
Intentional binding paradigm, response method, and readiness-potential extraction. **(A)** Schematic representation of the judgement error (JE), defined as the difference between the actual time of occurrence of an event (green circle, left: button press; red circle, right: tone) and its perceived time of occurrence (blue dots). In the Baseline conditions, the action and the tone occur in isolation; in the Operant conditions, the voluntary keypress produces the tone after a fixed 250 ms interval. The objective and perceived intervals are shown along the respective time axes (objective time for Baseline, subjective time for Operant). At the bottom, the action-binding (left arrow) and outcome-binding (right arrow) measures are illustrated, derived from the compression of the perceived interval relative to the objective interval. **(B)** The timing of perceived events was measured using the Libet clock paradigm for the assessment of IB. On each trial, participants verbally reported the perceived clock-hand position (blue dot) as an integer from 1 to 60, which the experimenter recorded. Green and red dots denote the actual time of the action and tone, respectively. **(C)** Statistical analyses on Readiness potentials (RP) were performed on the Cz electrode alone (outlined in red). For each trial, two RP components were extracted as the mean amplitude within two non-overlapping windows relative to the button press: the early RP (−2000 to −1000 ms) and the late RP (−500 to −50 ms).

#### Conditions and procedure

Four block types were presented, factorially crossing two judgement targets (Action vs Tone) with two agency contexts (Baseline vs Operant) (Fig. 2, Panel A). The Operant blocks and the Baseline Tone blocks comprised 4 blocks of 10 trials each; the Baseline Action condition comprised 2 blocks of 10 trials. Block order was pseudo-randomised across participants, with half completing the two Operant blocks first and the two Baseline blocks second, and the other half vice versa.

### Baseline Action

Participants pressed one of two functionally equivalent keys (Q or P) at a freely chosen moment after at least one full rotation of the clock hand. The use of two interchangeable keys, with instructions to alternate between them and avoid stereotyped sequences, was intended to maintain the voluntary nature of the movement and minimise automaticity. After the keypress, the clock hand continued rotating for 1500–2500 ms before disappearing. Participants then verbally reported the clock-hand position at the moment of pressing.

### Baseline Tone

The clock display was preceded by a 250-ms visual cue, followed at a random interval of 2500–6000 ms by a tone (500 ms duration). The clock hand continued rotating for 1500–2500 ms after the tone before disappearing. Participants then verbally reported the perceived clock-hand position at the moment of tone onset.

### Operant Action

Participants pressed one of the two keys (Q or P), and a tone was delivered 250 ms after the keypress. The clock hand continued rotating for 1500–2000 ms after the tone before disappearing. Participants then verbally reported the clock-hand position at the moment of keypress.

### Operant Tone

The procedure was identical to that of Operant Action, except that participants verbally reported the clock-hand position at the moment of tone onset.

#### EEG recordings

EEG was recorded with a 64-channel system (ActiCap; BrainAmp amplifier; BrainProducts, Gilching, Germany) positioned according to the international 10–20 system. FCz served as online reference and AFz as ground. Electrode impedances were kept below 10 kΩ. The signal was digitised at 1000 Hz and visualised online with BrainVision Recorder v1.21.0102. Event markers were transmitted from the stimulus-presentation computer to the amplifier via a TriggerBox (BrainProducts) and USB.

#### EEG preprocessing and Readiness Potential extraction

EEG data were processed offline with EEGLAB ^46^ and custom MATLAB scripts (MathWorks, Natick, MA). From the 64-channel acquisition cap, 17 fronto-central electrodes activated with conductive gel during the recording were retained as the active montage (Fz, F1, F2, FC1, FC2, FC3, FC4, Cz, C1, C2, C3, C4, CPz, CP1, CP2, T7, T8). The continuous signal was down-sampled to 250 Hz and band-pass filtered between 0.01 and 15 Hz using a zero-phase Hamming-windowed FIR filter ^47^. Data were then re-referenced to the common average of the active montage. Given the selected number of active scalp channels, Independent Component Analysis (ICA) was not applied. Artefact rejection was therefore performed at the trial level via automated amplitude thresholding: epochs in which any sample exceeded ±100 µV on any of the 17 electrodes were excluded from analysis. Subjects retaining fewer than 5 trials per condition after rejection were considered non-analysable. Continuous data were segmented into 3600-ms epochs spanning from −2600 to +1000 ms relative to the target event — the keypress in action conditions (Baseline Action, Operant Action) and the tone onset in tone conditions (Baseline Tone, Operant Tone). Baseline correction was applied using a pre-stimulus window from −2600 to −2500 ms, chosen to remain well outside the pre-movement preparatory activity window. RP amplitudes were quantified from the Cz electrode, conventionally used to index the RP because it overlies the vertex and captures the maximal amplitude of the medial frontocentral component generated bilaterally in the pre-supplementary and supplementary motor areas ^38^. For each trial, two RP components were extracted from Cz as the mean amplitude within two non-overlapping time windows: the early RP (−2000 to −1000 ms) and the late RP (−500 to −50 ms), both relative to the target event (Fig. 2, Panel C). From the initial 23 participants with both behavioural and EEG data, six were excluded from the EEG analyses: one for excessive artefact contamination after automated amplitude thresholding (>70% trials rejected); two for condition-dependent polarity inversion artefacts suggestive of unstable electrode contact, identified post hoc by visual inspection of subject-level QC plots; two for task interruption during the experimental session resulting in incomplete data for one or more conditions; and one for insufficient surviving trials per condition after automated artefact rejection in one or more of the four block types. The final EEG sample comprised 17 participants; the behavioural sample retained all 23.

#### Behavioural outcome measure

For each trial, the judgement error (JE) was calculated as the signed difference (in milliseconds) between the participant’s reported clock-hand position and the actual time of the relevant event (keypress in Action conditions; tone onset in Tone conditions). For the Operant Tone condition, where the tone followed the keypress by 250 ms, the reference position was adjusted to align the judgement to tone onset rather than to keypress onset. Differences exceeding ±30 clock units (half a clock rotation) were reflected by ±60 clock units to correct for wrap-around artefacts inherent to the circular clock display, and the resulting JE was converted to milliseconds using the clock calibration (1 unit = 42.7 ms). Negative JE values indicate anticipated awareness (event perceived before its true occurrence) while positive values indicate delayed awareness.

Median JE per condition per participant was the primary subject-level summary measure, chosen for its robustness to outliers relative to the mean ^48^ given the known variability of subjective timing judgements in Libet-clock paradigms ^49^. Action binding and outcome binding were defined as the Baseline–Operant shift in median JE:

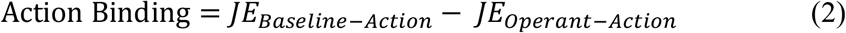

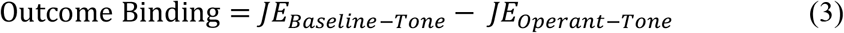

Under this convention, the canonical IB direction is opposite for the two binding measures: action binding is reflected by negative values (voluntary actions perceived as occurring later in the operant context than in baseline), whereas outcome binding is reflected by positive values (tones perceived as occurring earlier in the operant context than in baseline). Both directions reflect the temporal compression of the action–outcome interval that defines IB ^9^.

## Statistical Analysis

### Primary modelling framework

All trial-level analyses used linear mixed-effects models implemented in R (version 4.5.3) ^50^,with maximum-likelihood estimation and Satterthwaite degrees-of-freedom approximation. Fixed effects were tested via Type III ANOVA. The categorical FSS-7 classification (LowFSS vs HighFSS) was the primary analytic framework throughout, on three grounds: (i) the FSS-7 ≥ 5 threshold for high fatigue follows established use in the post-stroke fatigue literature ^44^ where it identifies stroke survivors with unambiguously high fatigue levels; (ii) the HighFSS distribution in the present cohort was tightly clustered (median 5.78, IQR 0.83), supporting its interpretation as a clinically homogeneous phenotype rather than a borderline group; (iii) continuous FSS-7 modelling was performed in parallel for confirmation of subject-level association with fatigue severity and is reported as a sensitivity analysis (Supplementary Tables S1, S3 and S4).

### Force Matching Task models

The primary mixed-effects model on SA included the target force level (State: 1, 2 or 3 N), fatigue group (LowFSS vs HighFSS) and their interaction as fixed effects, with a random intercept by block. State 20 N was excluded from the primary analysis because applied-force measurement at this force level was unreliable owing to apparatus saturation. The consequences of including it are reported as a validation analysis below. Subject-level analyses complemented the trial-level mixed-effects model with non-parametric tests of the same hypotheses: a Wilcoxon signed-rank test for SA ≠ 0 across all participants; Kruskal–Wallis comparisons of subject-mean SA, linear-fit gradient and intercept across HC, LowFSS and HighFSS; and a Spearman correlation between subject-level mean SA and FSS-7. Sensitivity analyses included (i) a conservative QC sample (≥ 75 % trial completion and exclusion of blocks with coefficient of variation > cohort mean + 3 SD); (ii) State 20 inclusion; (iii) continuous FSS-7 modelling.

### Intentional Binding behavioural models

The primary mixed-effects model on trial-level judgement error included Judgement (Action vs Tone), Agency (Baseline vs Operant), fatigue group (LowFSS vs HighFSS) and all their interactions as fixed effects, with a random intercept by participant. Where the focal three-way interaction was significant, estimated marginal means (EMMs) were extracted with the emmeans package using asymptotic degrees of freedom, and Baseline-vs-Operant contrasts were computed within each Judgement × FSS_group cell. As these contrasts were theoretically directional and pre-specified by the SAF account of PSF, P-values were reported without adjustment for multiple comparisons. Two sensitivity analyses tested the robustness of the focal interaction to mood: an additive model including HADS-Depression and HADS-Anxiety as fixed effects, and an extended model including HADS × FSS_group interactions.

## EEG (Readiness Potential) models

Trial-level EEG data were merged with behavioural data by ordinal trial rank within each subject × block-type cell, an alignment procedure necessary because trial indices in the EEG file referenced original event-related labels rather than behavioural trial number. Two primary mixed-effects models on trial-level mean amplitude at Cz were fitted, paralleling the structure of the behavioural analysis: one on the early RP window (−2000 to −1000 ms) and one on the late RP window (−500 to −50 ms), each with Agency (Baseline vs Operant), fatigue group (LowFSS vs HighFSS) and their interaction as fixed effects, and a random intercept by participant. EMMs and Baseline-vs-Operant contrasts within each FSS_group were extracted as for the behavioural models. Sensitivity analyses included HADS-Depression and HADS-Anxiety as additive covariates. Subject-level confirmation of the association with fatigue severity was provided by a Spearman correlation between FSS-7 and the subject-level RP modulation index (mean amplitude in Baseline − mean amplitude in Operant at the early RP window), with leave-one-out diagnostics across all 17 EEG participants.

### Robustness, equivalence and power analyses

The early-RP Agency x fatigue-group interaction was further probed with three convergent approaches suited to the modest sample: a permutation test with fatigue group shuffled at the subject level (5000 permutations), leave-one-out refitting of the interaction across all 17 participants, and a Bayesian mixed-effects estimate of the interaction using *brms* package ^51^ with weakly informative Normal(0, 5) priors, summarised as the posterior median and 95% credible interval. Estimated marginal contrasts are additionally reported as model-based standardised effects (Cohen’s d) with 95% confidence intervals. Predictive-postdictive uncoupling was tested directly by a partial Spearman correlation between the subject-level RP modulation index and outcome binding across the 17 patients with both measures, controlling for FSS-7. Cohort integration was assessed by paired two one-sided equivalence tests (TOST) on the 14 patients who completed both protocols. The equivalence bound for FSS-7 was pre-specified as half a standard deviation of the full IB cohort ^52^, in the absence of a validated minimal important difference for the seven-item scale. For the force-matching State x fatigue-group interaction, the minimum detectable effect at the achieved sample size was derived from the standard error of a one-degree-of-freedom (linear-State) parameterisation, used solely for this purpose and not as a significance test. Full results of these analyses are reported in Supplementary Table S5.

## Results

### Participants and cohort comparability

Two cohorts of chronic stroke survivors contributed to the present study: an Intentional Binding (IB) cohort (n = 23; 15 LowFSS, 8 HighFSS) and a Force Matching Task (FMT) cohort (n = 46; 31 LowFSS, 15 HighFSS), with 14 patients having completed both protocols at separate testing sessions (Table 1). The two cohorts were comparable on age (P = 0.93), FSS-7 distribution (P = 0.55), HADS-Depression (P = 0.52), HADS-Anxiety (P = 0.46) and hemiparesis distribution (P = 1.00). Within both cohorts, HADS-Depression scores were higher in HighFSS than in LowFSS patients, reaching statistical significance in the larger FMT cohort (median 7.0 versus 3.0; Mann–Whitney U, P = 0.005) and showing the same direction without reaching significance in the smaller IB cohort (median 7.5 versus 5.0; P = 0.38). To address potential mood confounding, the IB and EEG analyses were repeated with HADS-Depression and HADS-Anxiety as additive covariates (reported below). The block-level structure of the FMT recordings precluded individual-level covariate adjustment. For the 14 patients who completed both protocols, FSS-7 scores were stable across sessions (Spearman ρ = 0.72, P = 0.004), with 11 of 14 (79%) retaining the same fatigue-group classification and the three reclassified cases all lying close to the FSS-7 = 5 cut-off (Supplementary Table S2). This cross-session agreement supports the trait stability of fatigue severity despite the uncontrolled inter-session interval, while the within-cohort contrasts reported below depend only on each cohort’s own session classification.

Because each primary analysis was estimated within a single cohort, between-cohort differences do not enter the within-cohort fatigue contrasts, and cross-cohort comparability is reported here as contextual support. In the 14 patients assessed in both studies, fatigue severity did not differ between sessions (mean difference 0.33 FSS-7 points; paired t(13) = 0.99, P = 0.34) and lay within a pre-specified distribution-based bound of half a standard deviation (+/-0.87 points, derived from the full IB cohort), although formal equivalence was not reached at this sample size (two one-sided tests, P = 0.06); HADS-Depression and HADS-Anxiety likewise showed no session difference. Fatigue classification was stable across sessions (Spearman rho = 0.72; 11 of 14 concordant; median interval 3.7 years). The two datasets are therefore treated as convergent evidence from comparable cohorts rather than as a single integrated sample.

### Force Matching Task

#### Peripheral SA is preserved in PSF

The Force Matching Task probed peripheral, effector-level forward-model attenuation in 40 stroke survivors meeting the primary quality-control criterion (30 LowFSS, 10 HighFSS) and 24 healthy controls. Of the 46 recruited stroke survivors (Table 1), 42 provided analysable force-matching recordings, of which 40 satisfied the primary criterion (Supplementary Methods).

SA is detectable at group level. Across all 64 participants, matched force exceeded applied force in the great majority of trials (mean SA = 1.35 N, Wilcoxon signed-rank V = 1699, P < 0.0001), confirming that the basic forward-model attenuation mechanism is present in this combined sample at a level consistent with prior reports using the same apparatus ^5^.

No effect of fatigue on SA magnitude. Subject-level mean SA did not differ across the three groups (Kruskal–Wallis χ²(2) = 0.03, P = 0.99; Fig. 3, Panel A). The same null pattern was obtained for the secondary indices extracted from the per-subject linear fit relating matched to applied force: neither the gradient (χ²(2) = 0.65, P = 0.72) nor the intercept (χ²(2) = 0.12, P = 0.94) differed across HC, LowFSS and HighFSS. The group matched-versus-applied force relationship is shown in Supplementary Figure S3.

**Figure 3.**
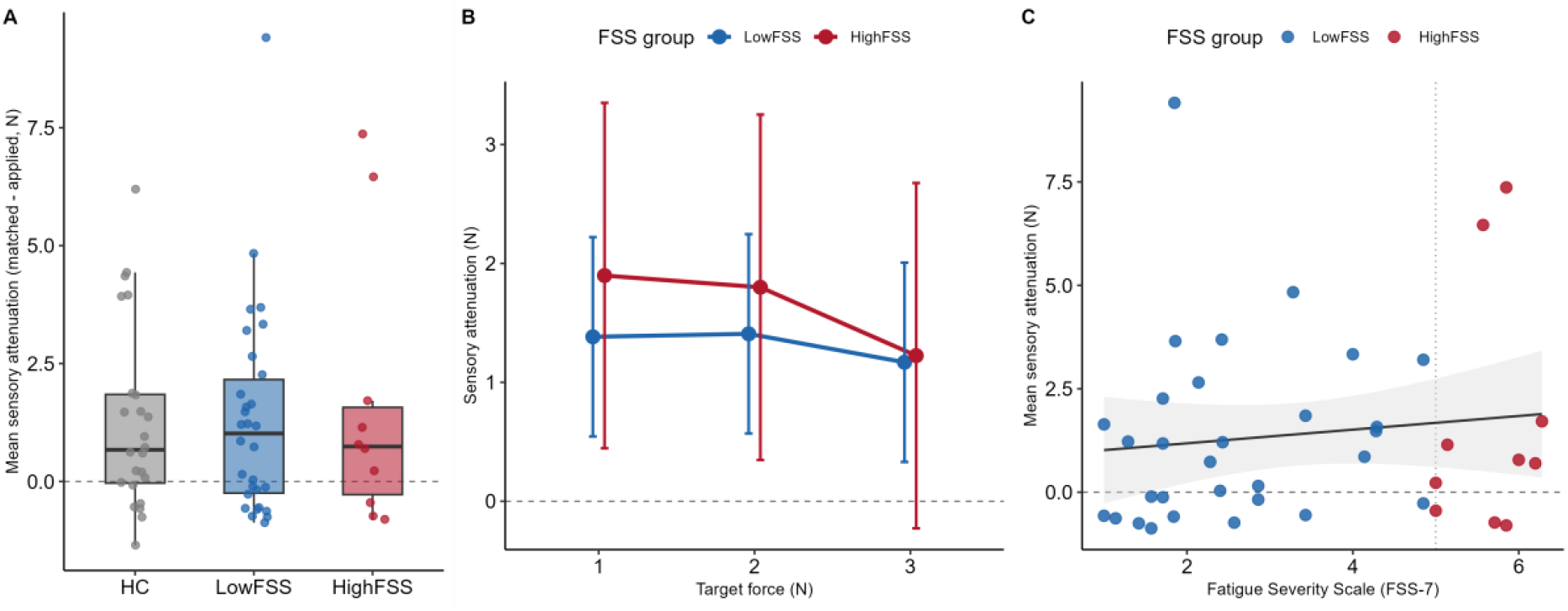
Peripheral SA is preserved across the fatigue range (FMT). Sensory attenuation (SA) is expressed as matched minus applied force (N); positive values indicate overestimation, the canonical signature of SA. **(A)** Subject-level mean SA for healthy controls (HC, grey), low-fatigue patients (LowFSS, blue) and high-fatigue patients (HighFSS, red); boxplots show median and interquartile range with individual participants overlaid. Groups did not differ (Kruskal–Wallis test reported in Results). **(B)** Estimated marginal means of SA as a function of target force (1, 2, 3 N) for LowFSS and HighFSS, from the primary trial-level mixed-effects model; error bars denote 95% confidence intervals. **(C)** Subject-level mean SA against continuous Fatigue Severity Scale score (FSS-7) in patients; the solid line and shaded band show the linear fit with 95% confidence interval, and the vertical dotted line marks the FSS-7 = 5 categorical cut-off. The dashed horizontal line at zero in all Panels indicates the absence of attenuation. Statistical values are reported in the Results and in Supplementary Table S1.

Extending the subject-level analysis with the primary trial-level mixed-effects model revealed the expected scaling of SA with target force (F(2, 4617) = 12.36, P < 0.0001) but no main effect of fatigue group (F(1, 40) = 0.14, P = 0.71; Fig. 3, Panel B; Supplementary Table S1, Panel A). The State × FSS_group interaction did not reach significance (F(2, 4617) = 2.88, P = 0.056), and three convergent analyses confirmed the absence of fatigue-dependent SA modulation: (i) all within-State LowFSS–HighFSS contrasts were non-significant (largest z = −0.60, P = 0.55; Fig. 3B, with broadly overlapping 95% confidence intervals); (ii) under a more conservative quality-control criterion (≥ 75% trial completion plus exclusion of blocks with coefficient of variation exceeding the cohort mean + 3 SD), the interaction remained non-significant (F(2, 4530) = 2.63, P = 0.07; Supplementary Table S1, Panel B); and (iii) the same model with continuous FSS-7 as predictor showed neither an interaction (F(2, 4617) = 1.67, P = 0.19) nor a main effect of FSS-7 on SA at trial level (F(1, 40) = 0.58, P = 0.45; Supplementary Table S1, Panel C), the latter confirmed at subject level by a negligible Spearman correlation between mean SA and FSS-7 (ρ = 0.15, P = 0.35; Fig. 3, Panel C). The central null of preserved SA across the FSS range therefore stands as the principal FMT experimental finding.

To express this preservation as a positive bound rather than a bare null, the minimum interaction effect detectable at the achieved sample size was estimated at approximately 0.28 N per force level (80% power), about one fifth of the mean attenuation (1.35 N). Any fatigue-related modulation of peripheral attenuation is therefore small relative to the attenuation itself (Supplementary Table S5, Panel C).

As a separate validation, we tested whether including the 20 N force level would alter the qualitative outcome. Including all four force levels produced a highly significant State × FSS_group interaction (F(3, 6185) = 20.28, P < 0.001), driven entirely by the 20 N condition, where applied-force measurement was unreliable owing to apparatus saturation (Supplementary Table S1, Panel D).

### Intentional Binding paradigm

#### Outcome binding is selectively amplified in fatigued patients

The Intentional Binding paradigm assessed implicit temporal compression of perceived intervals between voluntary actions and their tone outcomes in 23 chronic stroke survivors (15 LowFSS, 8 HighFSS).

#### Three-way interaction of Judgement, Agency and fatigue

The primary trial-level mixed-effects model on JE revealed a significant three-way Judgement × Agency × FSS_group interaction (F(1, 3185) = 7.39, P = 0.007; Fig. 4; Supplementary Table S3, Panel A). Lower-order effects included a robust main effect of Judgement (F(1, 3185) = 430.3, P < 0.0001), reflecting the expected difference between action-time and tone-time estimates, and significant Judgement × Agency (F(1, 3185) = 13.21, P < 0.001) and Judgement × FSS_group (F(1, 3185) = 14.86, P < 0.001) two-way interactions. The same three-way interaction was obtained when fatigue was modelled as a continuous FSS-7 predictor (F(1, 3185) = 7.99, P = 0.005; Supplementary Table S3, Panel B), confirming that the effect is not an artefact of the categorical split. The distribution of subject-level judgement error across the four conditions is shown in Supplementary Figure S1.

**Figure 4.**
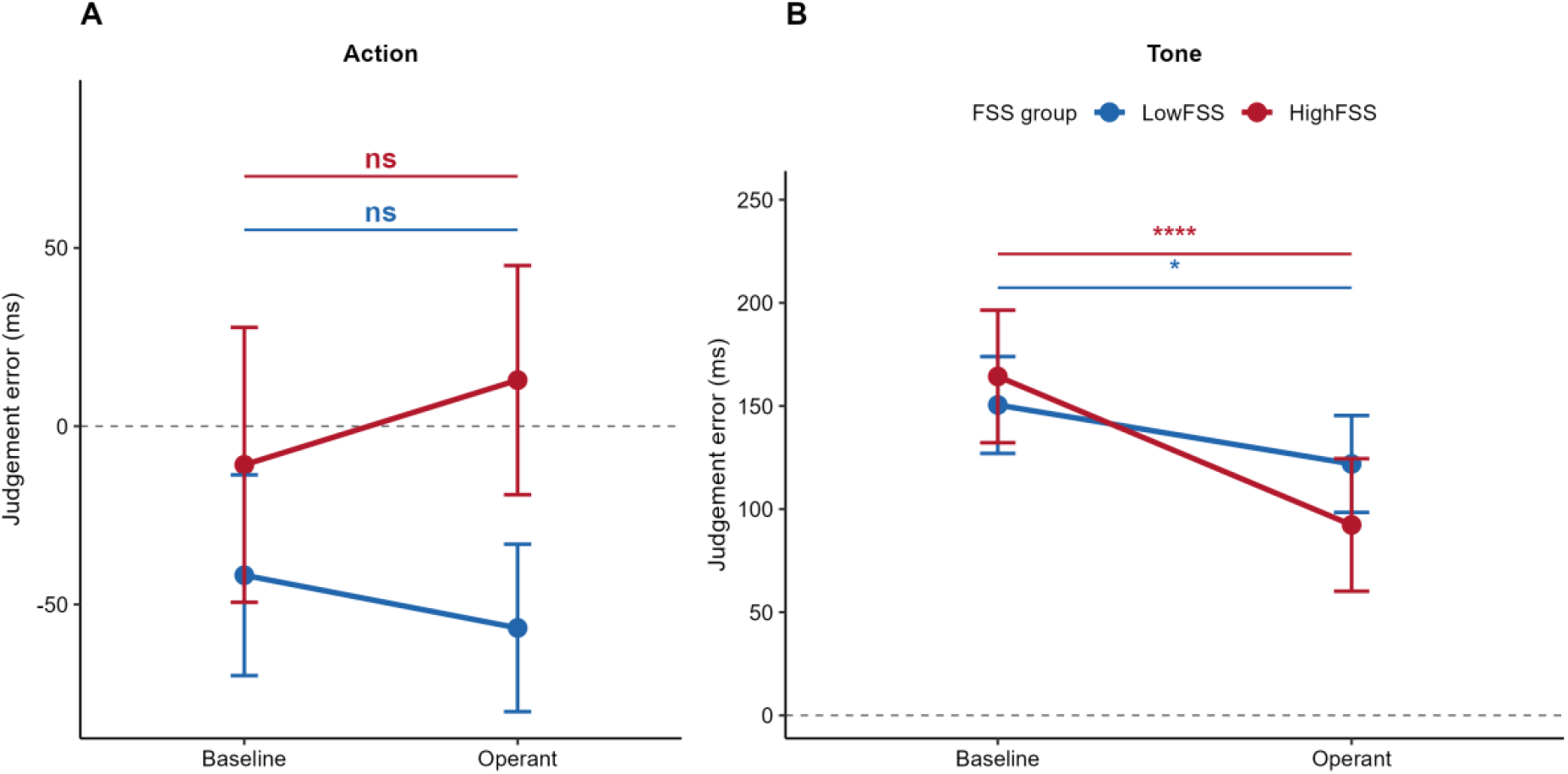
Outcome binding is selectively amplified in high-fatigue patients IB). Estimated marginal means of judgement error (ms) from the primary trial-level mixed-effects model, shown by Agency condition (Baseline vs Operant) for LowFSS (blue) and HighFSS (red); error bars denote 95% confidence intervals, and the dashed horizontal line marks zero judgement error. **(A)** Action judgement: the Baseline-to-Operant shift (action binding) did not reach significance in either group (ns). **(B)** Tone judgement: the Baseline-to-Operant shift (outcome binding) was significant in both groups and markedly larger in HighFSS than in LowFSS. Significance annotators denote within-group Baseline–Operant contrasts (* P < 0.05; **** P < 0.0001; ns, not significant); full estimates are reported in the Results. The fatigue-related modulation of IB is selectively localised to the postdictive (outcome) component, with action binding unmodulated by fatigue in both groups.

#### Outcome binding amplified in HighFSS

Estimated marginal means decomposed the three-way interaction along the theoretically critical contrast. In the Action judgement, the Baseline–Operant shift, indexing action binding, did not reach significance in either group (HighFSS: −23.7 ms, P = 0.21; LowFSS: 14.8 ms, P = 0.28; Fig. 4, Panel A). By contrast, in the Tone judgement, the Baseline–Operant shift indexing outcome binding was approximately 2.5 times larger in HighFSS than in LowFSS patients (Fig. 4, Panel B): HighFSS patients showed an outcome binding of 72.0 ms (z = 4.67, P < 0.0001), compared to 28.6 ms in LowFSS (z = 2.54, P = 0.01). The fatigue-related modulation of IB was therefore specifically localised to the postdictive (outcome) component, with action binding unmodulated in both groups. Subject-level distributions of the Action and Outcome binding indices, stratified by fatigue group, are shown in Supplementary Figure S2.

#### Robustness to mood

Adding HADS-Depression and HADS-Anxiety as additive covariates to the primary model left the three-way interaction essentially unchanged (F(1, 3185) = 7.38, P = 0.007; Supplementary Table S3, Panel C), and the same was true under a model including HADS × FSS_group interactions (F(1, 3185) = 7.38, P = 0.007; Supplementary Table S3, Panel D). HADS-Anxiety showed a small main effect on judgement error (F(1, 23) = 5.09, P = 0.03) but did not modulate the focal three-way interaction. HADS-Depression effects were not significant in any model. The HADS-adjusted estimated marginal means yielded numerically identical Baseline–Operant contrasts (HighFSS Tone: 72.0 ms, P < 0.0001; LowFSS Tone: 28.6 ms, P = 0.01), confirming that mood disturbance does not account for the fatigue-related modulation of outcome binding. A subject-level analysis of the binding indices, which does not detect the fatigue-related effect at the reduced power of the between-subject test (n = 23), is reported in Supplementary Table S3, Panel E.

### The early and late RPs fail to differentiate operant from baseline in HighFSS

Trial-level EEG data from a sub-sample of 17 patients (12 LowFSS, 5 HighFSS) were analysed to test whether the cortical preparatory complex, i.e., the descending predictive signal central to the SAF, is disrupted in fatigued patients. Grand-average RP at Cz across all four conditions are shown in Fig. 5A.

**Figure 5.**
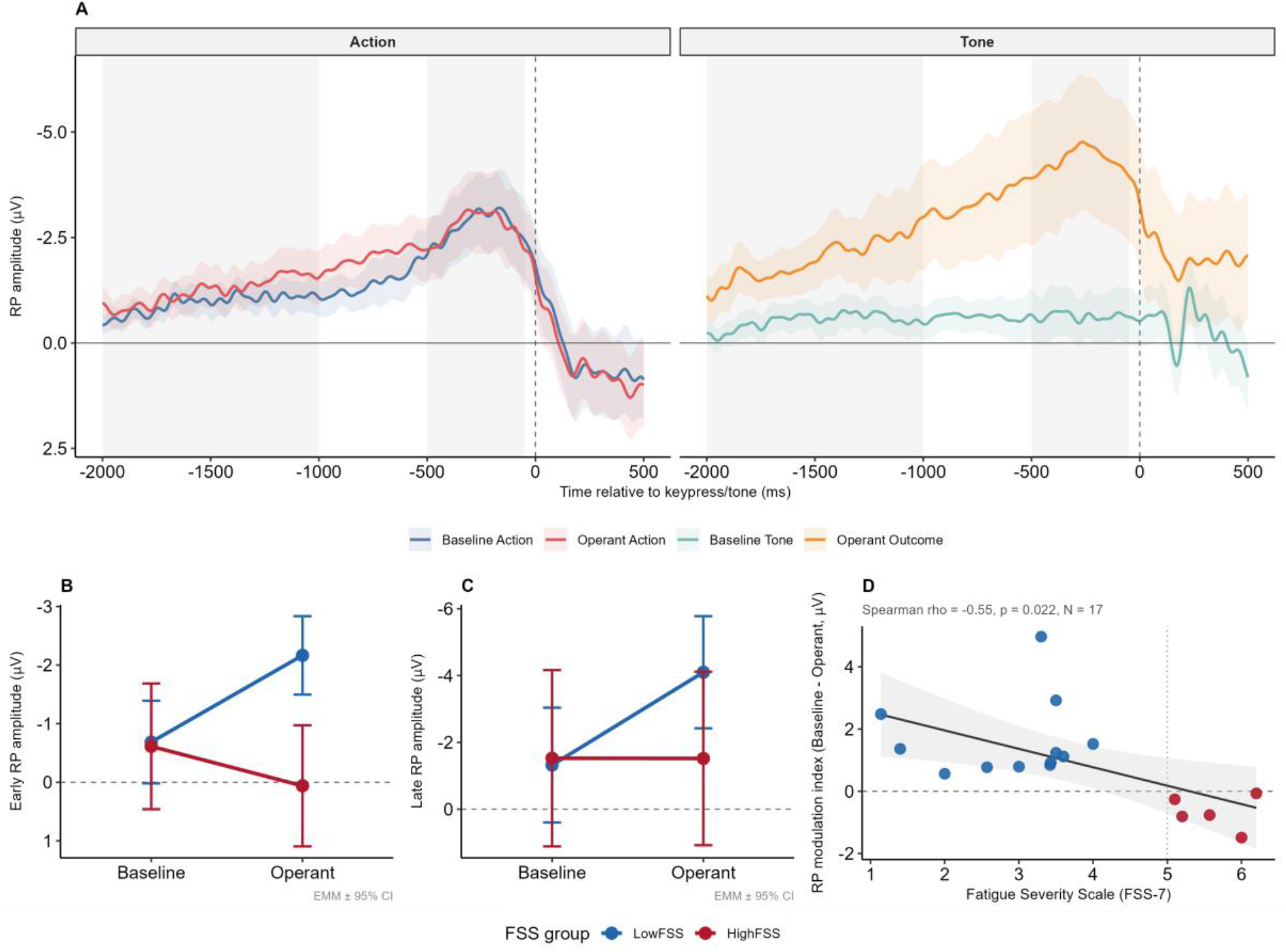
The readiness potential fails to differentiate operant from baseline in high-fatigue patients. Trial-level EEG from the 17-patient sub-sample (12 LowFSS, 5 HighFSS). **(A)** Grand-average RP at Cz across the four conditions — Baseline Action (blue), Operant Action (red), Baseline Tone (teal) and Operant Tone (orange) — time-locked to the keypress (Action conditions) or tone (Tone conditions) at 0 ms; shaded bands denote the standard error of the mean, and the shaded epoch marks the analysis windows. **(B)** Estimated marginal means of early RP amplitude (−2000 to −1000 ms) by Agency condition for LowFSS and HighFSS; LowFSS show the canonical operant-relative-to-baseline enhancement, absent in HighFSS. **(C)** Estimated marginal means of late RP amplitude (−500 to −50 ms), showing the same pattern. Error bars in (B) and (C) denote 95% confidence intervals; more negative amplitudes (plotted upward) indicate greater preparatory activity. **(D)** Subject-level RP modulation index (baseline minus operant mean amplitude at the early window) against FSS-7; the solid line and shaded band show the linear fit with 95% confidence interval, and the vertical dotted line marks the FSS-7 = 5 cut-off. Statistical values are reported in the Results and in Supplementary Table S4.

#### Early RP

The mixed-effects model on the mean amplitude at the early window (−2000 to −1000 ms) revealed a highly significant Agency × FSS_group interaction (*F* (1, 2043) = 13.72, *P* = 0.0002; Fig. 5B). Estimated marginal means showed that LowFSS patients exhibited the canonical enhancement of pre-motor preparatory activity in the operant relative to the baseline condition (Baseline − Operant = +1.48 μV, *z* = 4.61, *P* < 0.0001), whereas HighFSS patients showed no such differentiation (Baseline − Operant = −0.67 μV, *z* = −1.38, *P* = 0.17). The early RP enhancement that, in non-fatigued patients, signals agency-related pre-motor prediction was therefore absent in fatigued patients. Confirmation under continuous FSS modelling (F(1, 2039) = 9.64, P = 0.002) is reported in Supplementary Table S4, Panel E.

#### Late RP

A parallel analysis at the late window (−500 to-50 ms) revealed a comparable pattern (Agency × FSS_group: *F* (1, 2039) = 8.26, *P* = 0.004; Fig. 5C). LowFSS patients showed a robust Baseline–Operant enhancement of 2.78 μV (*z* = 5.21, *P* < 0.0001), whereas HighFSS patients showed essentially no modulation (Baseline−Operant = −0.007 μV, z = −0.01, P = 0.99). The agency-related differentiation of late pre-motor activity was therefore absent in fatigued patients. This pattern was confirmed under continuous FSS modelling (Agency × FSS-7: F (1, 2037) = 5.64, P = 0.018; Supplementary Table S4, Panel E).

#### Subject-level association with fatigue severity

To confirm the group-level findings at the level of individual variation, we computed for each subject the RP modulation index (mean baseline − mean operant amplitude at the early window) and correlated it with FSS-7. A negative correlation was observed (Spearman ρ =-0.55, *P* = 0.022; Fig. 5D), indicating that the attenuation of baseline-to-operant RP enhancement diminished progressively with fatigue severity across the full FSS-7 range. Leave-one-out diagnostics confirmed the correlation was not driven by individual outliers (range of leave-one-out ρ =-0.61 to-0.47, remaining significant at *P* < 0.05 in 12 of 17 iterations).

#### Robustness of the early-RP interaction

Given the modest EEG sample, the Agency x fatigue-group interaction was confirmed by three convergent approaches. A permutation test with fatigue group shuffled at the subject level supported the interaction (P = 0.0004, 5000 permutations); leave-one-out refitting left it significant in all 17 iterations (F range 9.12 to 14.14); and a Bayesian mixed-effects model placed the interaction at +2.12 µV (95% credible interval 0.98 to 3.25), with the posterior probability of the expected direction exceeding 0.99. Expressed as model-based standardised effects, the Baseline-Operant contrast was d = 0.25 (95% CI 0.14 to 0.35) in LowFSS and d =-0.11 (95% CI-0.27 to 0.05) in HighFSS. The interaction is thus robust. The directional conclusion for the high-fatigue subgroup, however, rests on five patients and is regarded as preliminary pending replication (Supplementary Table S5, Panel A).

#### Robustness to mood

Including HADS-Depression and HADS-Anxiety as additive covariates left both EEG interactions essentially unchanged (early RP: *F* (1, 2045) = 13.62, *P* = 0.0002; late RP: *F* (1, 2039) = 8.22, *P* = 0.004). Estimated marginal means under HADS adjustment yielded numerically equivalent Baseline–Operant contrasts at both windows (full tables in Supplementary Table S4).

## Discussion

Across three analytical blocks integrated from two independent cross-sectional studies on chronic stroke survivors, the present data delineate a coherent three-fold pattern that points to the central predictive layer of the sensory attenuation (SA) hierarchy as the locus of the agency-related impairment of post-stroke fatigue (PSF). Peripheral, effector-level forward-model attenuation, indexed by the force matching task (FMT), appears preserved across the Fatigue Severity Scale (FSS-7) range, a pattern that would distinguish PSF mechanistically from the peripheral SA failure phenotype observed in schizophrenia and functional movement disorders. Postdictive temporal integration, indexed by the outcome-binding component of the IB paradigm, is amplified in high-fatigue patients relative to low-fatigue patients. At the cortical level, preparatory activity indexed by the early and late readiness potential (RP) during voluntary action fails to differentiate operant from baseline conditions in high-fatigue patients, while showing the expected agency-related enhancement in low-fatigue patients. These results, jointly robust to mood adjustment, offer a first integrated empirical test of the sensory attenuation framework (SAF) account of PSF as an agency disorder, and are consistent with positioning it as an intermediate phenotype within the broader spectrum of agency-related disorders ^19,53^.

The FMT indicates that the basic forward-model attenuation mechanism remains operative across the FSS-7 range in our chronic stroke survivors. The pooled sample of 64 participants showed an unambiguous group-level effect, consistent in magnitude with prior reports using the same apparatus in healthy and clinical populations ^5^. Neither the subject-level mean nor the trial-level scaling of attenuation across target forces was modulated by fatigue. We interpret this preservation as informative rather than merely null, while acknowledging that the absence of an effect cannot by itself confirm fully intact peripheral processing. The FMT has documented sensitivity to peripheral SA failure in schizophrenia ^20^ and functional movement disorders ^24^, agency-related conditions from which the SAF predicts PSF should be mechanistically distinguishable. The same force-matching paradigm has also proved sensitive to graded change in Parkinson’s disease, where attenuation scales with disease severity and dopaminergic state ^54^, indicating that the task can register partial as well as near-abolished predictive deficits. Its insensitivity to fatigue here is therefore informative rather than a ceiling of the measure. The characteristic behaviour expected in these conditions is one of reduced overestimation, reflecting reduced or near-abolished descending predictive gain at the effector level. The absence of any comparable reduction across the FSS-7 range suggests that PSF does not entail an impairment of the peripheral forward model, i.e., the cancellation of predicted reafference at sensorimotor circuits proximal to muscle output, and would place the SAF-relevant disruption further upstream in the agency hierarchy, at the central preparatory layer addressed by the IB task.

The behavioural IB data reveal the first component of the central dissociation. IB, the implicit temporal index of the sense of agency introduced above, was selectively amplified in its postdictive component: high-fatigue patients showed an outcome binding approximately 2.5 times larger than low-fatigue patients, while the predictive component (action binding) was unmodulated by fatigue in both groups.

This pattern is interpretable within the cue-integration scheme ^2^, that argued that the sense of agency arises from the integration of two classes of cue: predictive cues, derived from the efference copy and available before the action, and postdictive cues, derived from the actual sensory feedback and available after it. The two are combined as a function of their relative reliability: a cue carrying more reliable information is weighted more heavily, and, conversely, when one cue is degraded, the system may rely proportionally more on the other. A comparable reliability-weighted reallocation of agency cues has been reported in other neurological contexts, with ageing increasing reliance on sensorimotor prediction ^55^ and sensory attenuation scaling with disease severity in Parkinson’s disease ^54^. When predictive cues are degraded (SAF-posited reduction in descending predictive gain supported directly by the RP analysis below), this scheme would predict a compensatory upweighting of postdictive inference, which could account for the observed inflated retrospective binding of sensory outcomes to the actions that caused them. The specific identity of the postdictive cue that may drive the amplification is theoretically in line with the SAF. Demanet and colleagues (2013) ^36^ manipulated physical effort independently of the primary IB task and found that IB increased under higher effort, concluding that effort sensations may be integrated as a non-specific cue to agency. SAF account incorporates this mechanism, positioning effort as the common denominator that links pathological fatigue to the agency-disorder spectrum: in fatigue, perceived effort is chronically elevated and disproportionate to objective motor demand, whereas in classical disorders of agency, it is altered differently, distorted in its attribution rather than amplified in its magnitude. Our findings are consistent with extending this principle to a pathological chronic-fatigue context: in high-fatigue patients, chronically elevated perceived effort, which the SAF attributes to reduced gain on descending motor predictions rather than to any peripheral attenuation deficit, may operate as a persistent and non-specific postdictive cue, amplifying the temporal compression of sensory outcomes toward the actions that caused them. An analogous amplification of retrospective, outcome-based binding alongside a degraded predictive component has been documented with the same paradigm in schizophrenia ^3^, providing an empirical precedent for this predictive–postdictive imbalance that we position more fully below.

One prediction ran counter to the data: action binding showed no significant Baseline–Operant contrast in either group, leaving the fatigue-related modulation selectively localised to the outcome component. While the limited statistical power of the sample to detect modest action-binding effects must be acknowledged, the selective localisation of the effect to the outcome component is also theoretically consistent with the proposal that effort signals act primarily as postdictive cues operating on outcome perception, rather than influencing the temporal estimation of the action itself. This pattern is consistent with evidence that action binding depends on efferent, prediction-based signals ^13^, whereas outcome binding is the more sensitive to inferential, postdictive cues about the action’s consequence ^12^.

The electrophysiological IB data complement behavioural findings by identifying a candidate neural substrate for the behavioural dissociation. In the operant condition, the keypress produces a tone outcome, whereas in the baseline condition, the same keypress carries no sensory consequence. The enhancement of preparatory activity in the operant relative to the baseline condition, therefore indexes the additional pre-motor processing engaged when an action is performed in order to produce a predictable sensory consequence, a candidate neural marker of agency-related, outcome-directed action. At the early window (−2000 to −1000 ms), low-fatigue patients displayed this canonical RP enhancement, reproducing the agency-related signature documented in healthy participants ^40^, whereas high-fatigue patients showed no such differentiation. At the late window (−500 to −50 ms) the same pattern emerged, but with greater amplitude.

The subject-level association with fatigue severity was consistent with the group-level effect at the individual level, converging toward the same interpretation: greater fatigue severity was associated with a progressively reduced operant-relative-to-baseline enhancement. The interpretive force of this finding rests on two converging lines of evidence that anchor the RP as a candidate neurophysiological index of agency-related computation, rather than of motor preparation exclusively. For example, transcranial magnetic stimulation of the pre-supplementary motor area, a principal cortical generator of the early RP ^38^, selectively reduces intentional binding, while stimulation of the sensorimotor cortex does not ^39^. Moreover, RP amplitude correlates within-subject with the magnitude of intentional binding in healthy participants ^40^. The early RP may therefore provide neural access to the descending predictive signal whose attenuation is the central SAF claim. Its absent agency-related modulation in high-fatigue patients is consistent with an impaired pre-motor predictive computation, although the small high-fatigue EEG subsample warrants caution in interpreting this absence. Abnormal premovement cortical potentials have been described across movement disorders, including reduced RP amplitude in Parkinson’s disease and an absent or variable potential in functional movement disorders ^56^. The present result differs from that literature in that it concerns not the amplitude or presence of the RP itself but the loss of its operant-versus-baseline modulation, an agency-specific contrast that isolates the predictive computation rather than motor preparation per se. This interpretation does not depend on interpreting the RP as a direct neural signature of intention. On the accumulator account, the pre-movement buildup partly reflects a leaky stochastic integration of subthreshold fluctuations crossing a decision threshold rather than a deterministic record of conscious planning ^57^. Crucially, the present inference rests not on the absolute RP but on the within-subject contrast between the baseline and operant conditions, which isolates the agency-specific modulation of preparatory activity regardless of how the baseline buildup itself is interpreted.

The convergent disruption of the late RP, a component anatomically distinct from the early one, originating predominantly in contralateral primary motor cortex and lateral premotor cortex ^38^, suggests that the deficit may not be confined to the genesis of the predictive signal alone but could extend to its downstream propagation through the preparatory motor system^58^ have shown that in healthy participants, the late RP becomes steeper when the sensory outcome of the action is anticipated, that is, the preparatory activity is shaped by the upcoming consequence. High-fatigue patients may fail to engage this outcome-anticipatory modulation, possibly because the upstream predictive computation that represents the action–outcome contingency is itself disrupted.

This electrophysiological signature does not stand in isolation but may supply the missing cognitive component of a symptom that has been progressively characterised at successively more proximal stages of the motor hierarchy. At resting state, high fatigue is associated with reduced resting corticomotor excitability and reduced excitability of the facilitatory inputs that drive motor cortex output ^33^. This hypoexcitability has a candidate network origin in the shift in interhemispheric inhibitory balance between homologous primary motor regions that predicts fatigue severity across independent samples ^34^. Closer to the motor act itself, this compromised system fails to reach the appropriate pre-movement state, showing attenuated corticospinal modulation during movement preparation ^35^. Each of these findings, however, characterises the motor system at the level of excitability and preparatory state. The absent agency-related RP modulation reported here may add the level at which that compromised preparatory state becomes behaviourally consequential. Therefore, we outlined a candidate trial-level cortical correlate of the failure to generate the descending predictive signal that, in an intact system, distinguishes an action performed to produce a sensory consequence from one that is not, thereby connecting the documented motor-excitability deficit of PSF to the alteration of agency computation.

The combined profile that emerges from these three analytical blocks (peripheral attenuation preserved, cortical predictive computation disrupted, postdictive integration amplified) is consistent with a system in which the predictive and postdictive components of agency become uncoupled by a single upstream impairment in descending predictive gain. In a physiologically calibrated agency system, the predictive and postdictive components are indeed coupled: the pre-action predictive signal constrains the retrospective integration of outcome cues, a coordination reflected in the within-subject correlation between RP amplitude and outcome binding documented in healthy participants ^40^. In high-fatigue patients this coordination appears to break down: with the predictive signal degraded, it may no longer constrain the postdictive integration, which would then operate with less calibration on the heightened salience of reafferent signals, including the effort signals whose magnitude defines the fatigue phenotype. It should be noted that this uncoupling is inferred from the co-occurrence of two group-level dissociations in the high-fatigue group, the amplified outcome binding and the attenuated agency-related RP modulation, rather than from a directly observed within-subject decorrelation of the two signals. Both were obtained in the same IB cohort, but outcome binding was available in all 23 patients whereas the readiness potential could be quantified only in the 17 with analysable EEG. The present design therefore does not test whether, within an individual patient, a smaller predictive modulation is accompanied by larger postdictive binding, the relationship that defines the healthy coupling. Thus, directly correlating subject-level RP modulation with outcome binding in the same patients would provide a more stringent test of the uncoupling hypothesis and remains an objective for future work. That the two readouts can dissociate at the level of measurement is itself expected. Recently, it has been argued that clock-based temporal estimates are retrospective perceptual constructions rather than direct readouts of motor preparatory activity ^49^. The predictive RP and the postdictive timing judgement can therefore be considered dissociable measures operating at different levels. Overall, the joint pattern observed across these dissociable measures in high-fatigue patients, the predictive RP modulation reduced, and the postdictive binding amplified, is consistent with a single upstream reduction in descending predictive precision giving rise to two distinct downstream consequences. This emerging profile appears mechanistically distinct from the agency-disorder pattern that defines schizophrenia and functional movement disorders, where the predictive-signal collapse is severe enough to produce a consistent breakdown of self-attribution ^20,23,24^. Notably, the same predictive and postdictive dissociation has been documented in schizophrenia using the IB paradigm ^3^. It has been shown that schizophrenic patients lack the predictive component of action awareness while exhibiting exaggerated retrospective binding, with the predictive deficit scaling with the severity of delusions and hallucinations. The PSF-related agency profile may share the computational form of this dissociation but not its severity or consequence: in PSF, descending predictive gain appears degraded but not abolished, self-attribution is categorically preserved, and the deficit manifests as a quantitative redistribution of the agency computation toward its postdictive component rather than as delusional reattribution. PSF may therefore represent not a disorder of diminished agency but a disorder of uncoupled agency, in which the predictive and postdictive components, normally coordinated, are dissociated by the same upstream impairment in descending predictive gain that, in the SAF account, is also proposed to underlie the heightened perceived effort that defines the clinical phenotype.

A fully within-subject, trial-level test of this coupling was not feasible here; we did, however, conduct the between-subject version in the 17 patients with both measures. The partial Spearman correlation between subject-level RP modulation and outcome binding, controlling for FSS-7, was negligible, as was the raw, group-confounded correlation (Supplementary Table S5, Panel B). This is consistent with an absence of residual predictive-postdictive coupling once fatigue is accounted for; however, the sample is powered only for large, so it cannot by itself establish uncoupling. Uncoupling is therefore advanced as a working hypothesis, supported rather than proven, by the present data.

The dissociation described above helps clarify the position that PSF may occupy within the broader family of agency-related disorders. At one extreme of this spectrum, schizophrenia and functional movement disorders exhibit a near-abolition of descending predictive gain: the failure cascades through the entire hierarchy, with peripheral SA reduced ^20,24^, self-attribution categorically compromised, and reafferent signals processed as if externally generated ^14,23^. At an intermediate position along this same spectrum, our data are consistent with a distinct configuration: peripheral SA preserved, cortical predictive computation disrupted but not abolished, and postdictive inference amplified rather than misattributed. The categorical experience of self-authorship is retained, while the deficit appears as a quantitative imbalance in the cue-integration scheme through which agency is computed. This configuration aligns with the phenotyping framework proposed for PSF ^53^. Arguing that pathological fatigue may be more productively conceptualised as a cluster of distinct disorders than as a unitary phenomenon, this framework identifies phenotyping as a precondition for both mechanistic understanding and intervention development. The signature emerging from the present investigation, may provide one such candidate mechanistic phenotype, defined at the computational rather than purely descriptive level. This computational phenotype may correspond to the motor-responsive, proprioceptive sub-group identified through corticospinal excitability and motor-cortex tDCS responsiveness ^32,53^, a correspondence whose interventional implications we consider below.

The framing of PSF as an intermediate agency-related phenotype may help reconcile three otherwise heterogeneous bodies of evidence: the phenomenology of effort elevation, disproportionate to objective motor demand and without delusional misattribution, that defines the PSF clinical picture, the well-documented but mechanistically unexplained pattern of cortical and corticospinal anomalies in high-fatigue patients and the empirical pattern reported here at the agency-computational level. All three would be accommodated by a single computational claim, that descending predictive gain is degraded but not abolished, together with the downstream consequences that follow from it.

Several methodological considerations qualify the present findings and bound the strength of the inferences they support.

The cross-sectional observational design constrains causal interpretation. The associations between fatigue severity, RP modulation, and outcome-binding amplification are consistent with the SAF prediction that descending predictive gain modulates both effort perception and agency computation, but the data themselves do not establish whether the predictive-gain deficit causes the fatigue, whether the fatigue compromises the predictive computation, or whether both reflect a common upstream factor (such as the network-level interhemispheric imbalance ^34^. Longitudinal designs tracking RP modulation alongside fatigue trajectory, or interventional designs manipulating cortical excitability, will be required to adjudicate these causal hypotheses. In the present work, we integrated the results from two independently conducted cross-sectional studies, while strengthening the interpretive framework by combining peripheral and central agency measures, which introduces between-study heterogeneity. The FMT and IB protocols were administered in separate sessions, with a convenience overlap of 14 patients participating in both. The cross-cohort comparisons establish demographic and clinical comparability but cannot eliminate residual sources of variance. Nevertheless, the within-cohort effects are unaffected by this consideration, and the convergence of the two within-cohort signatures around the SAF prediction strengthens rather than weakens the integrative conclusion. The intentional binding study did not include a healthy-control arm. Its two contrasts of interest, the within-subject agency contrast (operant versus baseline) and the between-group fatigue contrast (high-versus low-fatigue patients), were both estimated within the patient sample, with low-fatigue patients serving as the internal reference for the fatigue effect. The absence of a healthy-control arm therefore bears on the normative comparison between patients and healthy individuals rather than on these within-patient contrasts. Concerning the EEG sample (n = 17), modest numerosity reflects both the nature of the paradigm and the strict quality-control criteria adopted. In particular, the high-fatigue EEG subgroup was small (n = 5), so the absent RP differentiation in this group should be interpreted with corresponding caution. The leave-one-out diagnostics for the RP–FSS correlation indicated reasonably stable effects, but replication with larger samples and richer electrode coverage will be needed to consolidate the pattern. The choice to quantify the RP from the Cz electrode alone, although canonical in the RP literature ^38^ precludes topographic and source-localisation analyses. The decision not to apply ICA, driven by the limited number of active scalp channels, is consistent with related work on similarly constrained setups ^40,57^, but a fuller montage would enable more granular artefact decomposition in future studies. Finally, robustness analyses indicated that the findings based on the categorical fatigue distinction were more stable than their continuous-FSS counterparts. The categorical effects should therefore be regarded as the primary results, with continuous-FSS analyses reported as supporting evidence (Supplementary Tables S1, S3 and S4).

The small high-fatigue EEG subgroup (n = 5) noted above bounds the directional conclusion for that group. The interaction itself, however, survived permutation testing, Bayesian re-estimation and leave-one-out refitting, so this caution attaches to the magnitude of the high-fatigue effect rather than to its existence.

The present findings have potential implications for the development of mechanistically informed interventions for PSF. The localisation of the deficit at the cortical predictive layer of the agency hierarchy points to a specific neurophysiological substrate, descending predictive gain, indexed by the RP potential during voluntary action, as a candidate intervention target. Evidence support that motor cortex tDCS reduces PSF specifically by modifying the gain function of corticomotor networks, with the responsive sub-population corresponding to a proprioceptive phenotype defined by altered muscular sensory experience and the absence of mood disorder ^53^. The phenotype delineated by the present data, is a candidate computational definition of this sub-population. This would yield a testable prediction: a tDCS protocol calibrated to modulate cortical predictive gain might be expected to normalise both the agency-related RP modulation and the inflated outcome binding, in addition to reducing the fatigue itself.

Moreover, converging directions for future work emerge. Longitudinal studies tracking the joint trajectory of fatigue severity, RP modulation and binding from the subacute phase onward, when fatigue first emerges, would help establish the temporal precedence among these variables and constrain the causal architecture that a cross-sectional design cannot resolve. Replication with full-density EEG montages would clarify the relationship between early-and late-RP components ^58^. Sham-controlled neuromodulation with pre/post readouts of RP and binding would test the causal role of cortical predictive gain in the agency profile suggested here. While tDCS offers a scalable option, its diffuse focality limits inference about a specific generator, so focal pre-SMA stimulation would provide a more mechanistically specific causal test. Extension of the present paradigm to other pathological fatigue conditions, such as multiple sclerosis, in which fatigue has similarly been linked to functional network alteration rather than structural lesion load ^59^, would establish whether the agency-computational signature described here generalises across pathologies sharing the SAF mechanistic substrate or remains specific to the post-stroke aetiology.

Across behavioural, electrophysiological and theoretical levels, the present work delineates a coherent neuro computational profile of PSF where peripheral SA appears preserved, cortical predictive computation disrupted, and postdictive integration amplified. The dissociation among these three layers, consistent with joint consequences of a single upstream impairment in descending predictive gain, points to the cortical preparatory layer of the SA hierarchy as the node of the agency-related impairment of PSF. Moreover, it would distinguish PSF mechanistically from the peripheral-SA failure phenotype typical of schizophrenia and functional movement disorders and is consistent with positioning PSF as an intermediate phenotype within the broader spectrum of agency-related conditions. In this account, PSF may be characterised not as a disorder of diminished agency but as one of uncoupled agency, with the cortical predictive layer of the agency hierarchy emerging as a candidate level of both mechanistic disruption and potential therapeutic targeting.

## Data availability

The data underlying this article will be shared on reasonable request to the corresponding author.

## Acknowledgements

We thank Dr. S. Sporn for thoughtful discussions and assistance with the initial exploration and analysis of the Intentional Binding data. Generative AI (Claude, Anthropic) assisted with statistical script drafting and manuscript language editing; all outputs were reviewed and verified by the authors, who take full responsibility for the content.

## Supplementary material

Supplementary material is available at *Brain* online

## References

1. Haggard P. Sense of agency in the human brain. Nat Rev Neurosci. 2017;18(4):196–207. doi:10.1038/nrn.2017.14

2. Synofzik M, Vosgerau G, Voss M. The experience of agency: an interplay between prediction and postdiction. Front Psychol. 2013;4. doi:10.3389/fpsyg.2013.00127

3. Voss M, Moore J, Hauser M, Gallinat J, Heinz A, Haggard P. Altered awareness of action in schizophrenia: a specific deficit in predicting action consequences. Brain. 2010;133(10):3104–3112. doi:10.1093/brain/awq152

4. Blakemore SJ, Wolpert DM, Frith CD. Central cancellation of self-produced tickle sensation. Nat Neurosci. 1998;1(7):635–640. doi:10.1038/2870

5. Shergill SS, Bays PM, Frith CD, Wolpert DM. Two eyes for an eye: the neuroscience of force escalation. Science. 2003;301(5630):187. doi:10.1126/science.1085327

6. Press C, Kok P, Yon D. The Perceptual Prediction Paradox. Trends Cogn Sci. 2020;24(1):13–24. doi:10.1016/j.tics.2019.11.003

7. Wolpert DM, Ghahramani Z, Jordan MI. An internal model for sensorimotor integration. Science. 1995;269(5232):1880-1882. doi:10.1126/science.7569931

8. Gu J, Buidze T, Zhao K, Gläscher J, Fu X. The neural network of sensory attenuation: A neuroimaging meta-analysis. Psychon Bull Rev. 2025;32(1):31–51. doi:10.3758/s13423-024-02532-1

9. Haggard P, Clark S, Kalogeras J. Voluntary action and conscious awareness. Nat Neurosci. 2002;5(4):382–385. doi:10.1038/nn827

10. Moore JW, Obhi SS. Intentional binding and the sense of agency: A review. Consciousness and Cognition. 2012;21(1):546–561. doi:10.1016/j.concog.2011.12.002

11. Libet B, Gleason CA, Wright EW, Pearl DK. Time of conscious intention to act in relation to onset of cerebral activity (readiness-potential). The unconscious initiation of a freely voluntary act. Brain. 1983;106 (Pt 3):623–642. doi:10.1093/brain/106.3.623

12. Moore JW, Wegner DM, Haggard P. Modulating the sense of agency with external cues. Consciousness and Cognition. 2009;18(4):1056–1064. doi:10.1016/j.concog.2009.05.004

13. Engbert K, Wohlschläger A, Haggard P. Who is causing what? The sense of agency is relational and efferent-triggered. Cognition. 2008;107(2):693–704. doi:10.1016/j.cognition.2007.07.021

14. Brown H, Adams RA, Parees I, Edwards M, Friston K. Active inference, sensory attenuation and illusions. Cogn Process. 2013;14(4):411–427. doi:10.1007/s10339-013-0571-3

15. Friston KJ, Daunizeau J, Kilner J, Kiebel SJ. Action and behavior: a free-energy formulation. Biol Cybern. 2010;102(3):227–260. doi:10.1007/s00422-010-0364-z

16. Moore J, Haggard P. Awareness of action: Inference and prediction. Consciousness and Cognition. 2008;17(1):136–144. doi:10.1016/j.concog.2006.12.004

17. Marcora S. Perception of effort during exercise is independent of afferent feedback from skeletal muscles, heart, and lungs. J Appl Physiol (1985). 2009;106(6):2060-2062. doi:10.1152/japplphysiol.90378.2008

18. de Morree HM, Klein C, Marcora SM. Perception of effort reflects central motor command during movement execution. Psychophysiology. 2012;49(9):1242–1253. doi:10.1111/j.1469-8986.2012.01399.x

19. Kuppuswamy A. The fatigue conundrum. Brain. 2017;140(8):2240–2245. doi:10.1093/brain/awx153

20. Shergill SS, Samson G, Bays PM, Frith CD, Wolpert DM. Evidence for sensory prediction deficits in schizophrenia. Am J Psychiatry. 2005;162(12):2384–2386. doi:10.1176/appi.ajp.162.12.2384

21. Lafargue G, Franck N, Sirigu A. Sense of motor effort in patients with schizophrenia. Cortex. 2006;42(5):711–719. doi:10.1016/s0010-9452(08)70409-x

22. Lafargue G, Franck N. Effort awareness and sense of volition in schizophrenia. Conscious Cogn. 2009;18(1):277–289. doi:10.1016/j.concog.2008.05.004

23. Frith CD, Done DJ. Experiences of alien control in schizophrenia reflect a disorder in the central monitoring of action. Psychol Med. 1989;19(2):359–363. doi:10.1017/s003329170001240x

24. Pareés I, Brown H, Nuruki A, et al. Loss of sensory attenuation in patients with functional (psychogenic) movement disorders. Brain. 2014;137(11):2916–2921. doi:10.1093/brain/awu237

25. Fletcher PC, Frith CD. Perceiving is believing: a Bayesian approach to explaining the positive symptoms of schizophrenia. Nat Rev Neurosci. 2009;10(1):48–58. doi:10.1038/nrn2536

26. Chaudhuri A, Behan PO. Fatigue in neurological disorders. Lancet. 2004;363(9413):978–988. doi:10.1016/S0140-6736(04)15794-2

27. De Doncker W, Dantzer R, Ormstad H, Kuppuswamy A. Mechanisms of poststroke fatigue. J Neurol Neurosurg Psychiatry. 2018;89(3):287–293. doi:10.1136/jnnp-2017-316007

28. Kuppuswamy A. The Neurobiology of Pathological Fatigue: New Models, New Questions. Neuroscientist. 2022;28(3):238–253. doi:10.1177/1073858420985447

29. Krupp LB, LaRocca NG, Muir-Nash J, Steinberg AD. The Fatigue Severity Scale: Application to Patients With Multiple Sclerosis and Systemic Lupus Erythematosus. Arch Neurol. 1989;46(10):1121–1123. doi:10.1001/archneur.1989.00520460115022

30. Johansson S, Kottorp A, Lee KA, Gay CL, Lerdal A. Can the Fatigue Severity Scale 7-item version be used across different patient populations as a generic fatigue measure – a comparative study using a Rasch model approach. Health Qual Life Outcomes. 2014;12:24. doi:10.1186/1477-7525-12-24

31. Lerdal A, Kottorp A. Psychometric properties of the Fatigue Severity Scale-Rasch analyses of individual responses in a Norwegian stroke cohort. Int J Nurs Stud. 2011;48(10):1258–1265. doi:10.1016/j.ijnurstu.2011.02.019

32. De Doncker W, Charles L, Ondobaka S, Kuppuswamy A. Exploring the relationship between effort perception and poststroke fatigue. Neurology. 2020;95(24):e3321–e3330. doi:10.1212/WNL.0000000000010985

33. Kuppuswamy A, Clark EV, Turner IF, Rothwell JC, Ward NS. Post-stroke fatigue: a deficit in corticomotor excitability? Brain. 2015;138(Pt 1):136–148. doi:10.1093/brain/awu306

34. Ondobaka S, De Doncker W, Ward N, Kuppuswamy A. Neural effective connectivity explains subjective fatigue in stroke. Brain. 2022;145(1):285–294. doi:10.1093/brain/awab287

35. De Doncker W, Brown KE, Kuppuswamy A. Influence of post-stroke fatigue on reaction times and corticospinal excitability during movement preparation. Clin Neurophysiol. 2021;132(1):191–199. doi:10.1016/j.clinph.2020.11.012

36. Demanet J, Muhle-Karbe PS, Lynn MT, Blotenberg I, Brass M. Power to the will: how exerting physical effort boosts the sense of agency. Cognition. 2013;129(3):574–578. doi:10.1016/j.cognition.2013.08.020

37. Kornhuber HH, Deecke L. Changes in the brain potential in voluntary movements and passive movements in man: readiness potential and reafferent potentials. Pflugers Arch Gesamte Physiol Menschen Tiere. 1965;284:1–17.

38. Shibasaki H, Hallett M. What is the Bereitschaftspotential? Clin Neurophysiol. 2006;117(11):2341–2356. doi:10.1016/j.clinph.2006.04.025

39. Moore JW, Ruge D, Wenke D, Rothwell J, Haggard P. Disrupting the experience of control in the human brain: pre-supplementary motor area contributes to the sense of agency. Proc Biol Sci. 2010;277(1693):2503–2509. doi:10.1098/rspb.2010.0404

40. Jo HG, Wittmann M, Hinterberger T, Schmidt S. The readiness potential reflects intentional binding. Front Hum Neurosci. 2014;8:421. doi:10.3389/fnhum.2014.00421

41. Stenner MP, Haggard P. Voluntary or involuntary? A neurophysiologic approach to functional movement disorders. Handb Clin Neurol. 2016;139:121–129. doi:10.1016/B978-0-12-801772-2.00011-4

42. Zapparoli L, Seghezzi S, Devoto F, et al. Altered sense of agency in Gilles de la Tourette syndrome: behavioural, clinical and functional magnetic resonance imaging findings. Brain Commun. 2020;2(2):fcaa204. doi:10.1093/braincomms/fcaa204

43. Snaith RP. The Hospital Anxiety And Depression Scale. Health Qual Life Outcomes. 2003;1:29. doi:10.1186/1477-7525-1-29

44. Kuppuswamy A, Clark EV, Sandhu KS, Rothwell JC, Ward NS. Post-stroke fatigue: a problem of altered corticomotor control? J Neurol Neurosurg Psychiatry. 2015;86(8):902–904. doi:10.1136/jnnp-2015-310431

45. Brainard DH. The Psychophysics Toolbox. Spat Vis. 1997;10(4):433–436.

46. Delorme A, Makeig S. EEGLAB: an open source toolbox for analysis of single-trial EEG dynamics including independent component analysis. J Neurosci Methods. 2004;134(1):9–21. doi:10.1016/j.jneumeth.2003.10.009

47. Widmann A, Schröger E, Maess B. Digital filter design for electrophysiological data--a practical approach. J Neurosci Methods. 2015;250:34–46. doi:10.1016/j.jneumeth.2014.08.002

48. Ratcliff R. Methods for dealing with reaction time outliers. Psychol Bull. 1993;114(3):510–532. doi:10.1037/0033-2909.114.3.510

49. Triggiani AI, Kreiman G, Lewis C, et al. What is the intention to move and when does it occur? Neuroscience & Biobehavioral Reviews. 2023;151:105199. doi:10.1016/j.neubiorev.2023.105199

50. Kuznetsova A, Brockhoff PB, Christensen RHB. lmerTest Package: Tests in Linear Mixed Effects Models. Journal of Statistical Software. 2017;82:1–26. doi:10.18637/jss.v082.i13

51. Bürkner PC. brms: An R Package for Bayesian Multilevel Models Using Stan. Journal of Statistical Software. 2017;80:1–28. doi:10.18637/jss.v080.i01

52. Norman GR, Sloan JA, Wyrwich KW. Interpretation of changes in health-related quality of life: the remarkable universality of half a standard deviation. Med Care. 2003;41(5):582–592. doi:10.1097/01.MLR.0000062554.74615.4C

53. Kuppuswamy A. Post-stroke fatigue – a multidimensional problem or a cluster of disorders? A case for phenotyping post-stroke fatigue. The Journal of Physiology. 2025;603(3):759–772. doi:10.1113/JP285900

54. Wolpe N, Zhang J, Nombela C, Ingram JN, Wolpert DM, Rowe JB. Sensory attenuation in Parkinson’s disease is related to disease severity and dopamine dose. Sci Rep. 2018;8(1):15643. doi:10.1038/s41598-018-33678-3

55. Wolpe N, Ingram JN, Tsvetanov KA, et al. Ageing increases reliance on sensorimotor prediction through structural and functional differences in frontostriatal circuits. Nat Commun. 2016;7(1):13034. doi:10.1038/ncomms13034

56. Colebatch JG. Bereitschaftspotential and movement-related potentials: origin, significance, and application in disorders of human movement. Mov Disord. 2007;22(5):601–610. doi:10.1002/mds.21323

57. Schurger A, Sitt JD, Dehaene S. An accumulator model for spontaneous neural activity prior to self-initiated movement. Proc Natl Acad Sci U S A. 2012;109(42):E2904–2913. doi:10.1073/pnas.1210467109

58. Kong G, Barlerin B, Desoche C, et al. Spatial Distance and Temporal Attentional Focus Modulate Voluntary Action Preparation and Awareness. Psychophysiology. 2026;63(3):e70280. doi:10.1111/psyp.70280

59. Bertoli M, Tecchio F. Fatigue in multiple sclerosis: Does the functional or structural damage prevail? Mult Scler. 2020;26(14):1809–1815. doi:10.1177/1352458520912175

