## Supplementary files for "Post-stroke fatigue as a disorder of uncoupled agency: an exploration of predictive and postdictive mechanisms"

### Supplementary Methods

#### 1. Force matching task: apparatus calibration

The force sensor used in the force matching task (FMT) acquired data at 2,500 Hz and produced output in raw voltage units. The device-specific calibration factor was 0.0197 V = 1 N, equivalent to a multiplicative factor of 50.76 for converting voltage to Newtons. The calibration was verified post hoc by extracting the median applied force per target State (1 N, 2 N, 3 N) and confirming that calibrated values matched the experimental target. Across the pooled sample, post-calibration mean sensory attenuation was 1.35 N, consistent in magnitude with prior reports using the same apparatus[1].

#### 2. Force matching task: quality control criteria

**Primary criterion.** Subjects were retained if they completed at least 50% of trials and provided a usable force matching response on each retained trial. From the initial FMT cohort (n = 46; 31 LowFSS, 15 HighFSS), this criterion retained 40 patients (30 LowFSS, 10 HighFSS) for primary analysis. Exclusions comprised subjects with extensive missing-trial patterns or non-monotonic matched-force responses across target levels. Twenty-four healthy controls (HC) completed the same protocol and were retained under the identical criterion; healthy-control blocks carry no FSS or demographic labelling and were used only for the group-level comparisons (Wilcoxon signed-rank, Kruskal–Wallis) and Figure S3.

**Conservative criterion.** As a sensitivity analysis, a more conservative quality check (QC) was applied: subjects were retained if they completed at least 75% of trials, and blocks with within-block coefficient of variation (CoV) exceeding the cohort mean + 3 SD were excluded. This conservative criterion retained 39 patients.

**State 20 N exclusion.** The 20 N force level was excluded from the primary analysis because applied-force measurement at this force level was unreliable owing to apparatus saturation: the force sensor approaches its dynamic range ceiling, resulting in compressed and non-linear voltage-to-force conversion that distorts the matched-applied relationship. The empirical consequence of including State 20 (a highly significant State × FSS group interaction driven entirely by the saturated condition ) is reported in Table S1 (Panel D).

#### 3. Cross-cohort matching tests

Comparability between the IB and FMT cohorts was tested for each demographic and clinical variable (age, FSS-7, HADS-Depression, HADS-Anxiety, hemisphere distribution) using the Mann–Whitney U test for continuous variables and Fisher's exact test for hemisphere distribution. All cross-cohort comparisons were non-significant at conventional alpha ($P > 0.4$ for all variables;

see Table 1 in the main manuscript). Within-cohort comparisons of LowFSS vs HighFSS patients confirmed the expected FSS-7 group separation in both cohorts ($P < 0.001$) and consistently showed HADS-Depression scores higher in HighFSS patients across both cohorts — reaching significance in the larger FMT cohort ($P = 0.005$) and showing the same directional pattern without reaching significance in the smaller IB cohort ($P = 0.38$). Cross-session agreement of FSS-7 between the IB and FMT sessions, computed in the 14 overlap participants, was good (Spearman $\rho = 0.72$, $P = 0.004$ and 11/14 retained the same fatigue-group classification; see Table S2).

The equivalence bound for this cross-session comparison was pre-specified as half a standard deviation of the full 23-patient IB cohort (SD = 1.74; bound +/-0.87 FSS-7 points), following the distribution-based convention of Norman, Sloan and Wyrwich [2], and adopted in the absence of a validated minimal important difference for the seven-item Fatigue Severity Scale. For comparison, the same test using a standard deviation computed within the 14-patient overlap sample (SD = 1.93; bound +/-0.97) did reach formal equivalence ($P = 0.04$), illustrating that the outcome depends on the reference standard deviation at this sample size; the cohort-derived bound (+/-0.87) is reported as the primary, non-circular estimate. HADS-Depression and HADS-Anxiety showed no session difference under the equivalent procedure. Equivalence bounds were fixed a priori.

### 4. Intentional Binding: EEG preprocessing additional details

**Active electrode montage.** Recordings were performed using a 64-channel BrainAmp cap. Conductive gel was applied to a pre-defined fronto-central subset of 17 electrodes that constituted the active montage for analysis: Fz, F1, F2, FC1, FC2, FC3, FC4, Cz, C1, C2, C3, C4, CPz, CP1, CP2, T7, T8. Remaining channels were not gel-coupled and contained no analysable signal. RP amplitudes for the statistical analyses were quantified from Cz alone (see main Methods).

**Event markers and trigger summation.** Event markers used to identify trial epochs followed a standard scheme: S 13 = Baseline Action keypress; S 64/S 65 = Baseline Tone onset; S 113 = Operant Action keypress; S 163 = Operant Outcome keypress (with the tone delivered 250 ms after the keypress in the Operant Tone condition). For subject 12, marker summation with a response-box trigger (S 128) shifted all marker values by +128; the corresponding shifted markers were correctly identified during preprocessing. The Operant Outcome marker S 163 was not affected by the summation because the tone arrived after the keypress and was not coincident with the response-box trigger.

**Task restart handling.** Two participants experienced task interruption during the experimental session, producing two recording sessions within the same EEG file. The latency of the second experiment-start marker was identified for each, and all events post-restart were excluded from analysis. Both participants had insufficient surviving trials per condition and were excluded from the final EEG dataset (see exclusion summary, main Methods).

**5. Statistical software and session information**

All mixed-effects analyses were performed in R using the lme4 package [3] and the lmerTest package [4](Kuznetsova, Brockhoff & Christensen, 2017), with Type III ANOVA tables computed via Satterthwaite's approximation for the denominator degrees of freedom. Estimated marginal means and Baseline-vs-Operant contrasts were extracted with the emmeans package[5] using asymptotic degrees of freedom. Models were fitted using maximum likelihood (ML) for nested-model comparisons and REML for parameter estimation. Spearman correlations and Mann–Whitney / Kruskal–Wallis tests were computed using base R.

**6. Direct test of predictive-postdictive uncoupling**

Because a fully within-subject, trial-level test of the RP-binding coupling documented in healthy participants[6] was not feasible with the present design, the feasible between-subject analogue was conducted in the 17 patients with both readiness-potential and binding measures. For each patient, the subject-level RP modulation index (Baseline minus Operant, early window) and the outcome-binding index (Baseline minus Operant, Tone judgement) were correlated using Spearman's rho, both raw and partial (controlling for FSS-7; ppcor package [7]). Because the subject-level binding indices carry substantial noise (interquartile ranges up to approximately 160 ms across fatigue groups; Table S3, Panel E), and because this between-subject test is a proxy for, rather than equivalent to, the within-subject/trial-level coupling reported in healthy participants, the analysis is powered only for large associations (rho of approximately 0.6 or greater at $n = 17$). The result is reported in Table S5b and interpreted in the main Discussion as consistent with, but not establishing, predictive-postdictive uncoupling.

**7. Robustness analyses for the early readiness-potential interaction**

Given the modest EEG sample ($n = 17$; HighFSS $n = 5$), the early-RP Agency x FSS_group interaction was subjected to three convergent robustness analyses. First, a permutation test shuffled the FSS_group label at the subject level (preserving within-subject dependence across Agency conditions) across 5000 permutations, refitting the interaction F-statistic each time. Second, leave-one-out refitting excluded each of the 17 participants in turn and recomputed the interaction. Third, a Bayesian mixed-effects model with the same fixed- and random-effects structure was fitted using the brms package[8] (rstan backend; weakly informative Normal(0, 5) priors on fixed effects; 4 chains, 4000 iterations per chain, seed = 1), summarised as the posterior median, 95% credible interval and the posterior probability of the expected-direction coefficient. Convergence diagnostics (R-hat, effective sample size) were not extracted for the present summary and are available on request. Results are reported in Table S5a.

**8. Minimum detectable effect and equivalence testing for the force-matching interaction**

To express the preserved sensory-attenuation result as a positive bound rather than a bare null, two complementary analyses were performed on the State x FSS_group interaction (Table S1, Panel A; P = 0.056). First, a one-degree-of-freedom linear-State parameterisation was fitted solely to obtain the standard error of the interaction term (not as a significance test); the minimum detectable effect (MDE) at a given power was then derived analytically as MDE = (z(1-alpha/2) + z(power)) x SE, giving 2.80 x SE for 80% power and 3.24 x SE for 90% power. Second, two one-sided equivalence tests (TOST; TOSTER package [9,10]) were performed on the subject-level SA indices (mean SA, gradient, intercept) between LowFSS and HighFSS patients, using a pre-specified bound of +/-0.5 standard deviations. This bound was adopted as a conventional default in the absence of a natural raw-unit anchor for these indices; the TOSTER package itself flags that standardised-mean-difference bounds can give biased equivalence estimates in small samples, a caveat that applies to the present HighFSS subgroup (n = 10) and is noted here for transparency. Full results are reported in Table S5c.

### Supplementary Tables

#### Table S1. Force matching task: sensitivity analyses

Type III ANOVA tables (Satterthwaite degrees of freedom) for the trial-level mixed-effects model of sensory attenuation as a function of target force level and fatigue group, with a random intercept by block. Panel A reports the primary analysis (n = 40 patients, States 1–3 N); Panel B reports the conservative quality-control sample; Panel C reports the same primary model with continuous FSS-7 in place of categorical FSS_group; Panel D reports the consequence of including the unreliable State 20 N condition.

***Panel A. Primary model (n = 40; States 1–3 N)***

| Effect | Num df | Den df | *F* | *P* |
|---|---|---|---|---|
| State (target force) | 2 | 4617 | 12.36 | <0.0001 |
| FSS group | 1 | 40.0 | 0.14 | 0.710 |
| State × FSS group | 2 | 4617 | 2.88 | 0.056 |

***Panel B. Conservative QC sample (≥ 75% trial completion + CoV outlier exclusion)***

| Effect | Num df | Den df | *F* | *P* |
|---|---|---|---|---|
| State (target force) | 2 | 4530 | 11.55 | <0.0001 |
| FSS group | 1 | 39.0 | 0.45 | 0.505 |
| State × FSS group | 2 | 4530 | 2.63 | 0.072 |

***Panel C. Continuous FSS-7 modelling***

| Effect | Num df | Den df | *F* | *P* |
|---|---|---|---|---|
| State (target force) | 2 | 4617 | 9.66 | <0.0001 |
| FSS-7 (continuous) | 1 | 40.0 | 0.58 | 0.450 |
| State × FSS-7 | 2 | 4617 | 1.67 | 0.189 |

***Panel D. State 20 N inclusion (all four force levels)***

| Effect | Num df | Den df | *F* | *P* |
|---|---|---|---|---|
| State (target force) | 3 | 6185 | 610.18 | <0.001 |
| FSS group | 1 | 40.0 | 0.68 | 0.413 |
| State × FSS group | 3 | 6185 | 20.28 | <0.0001 |

The State × FSS group interaction did not reach significance in the primary model (Panel A, P = 0.056) and remained non-significant under the conservative quality-control sample (Panel B, P = 0.072) and under continuous FSS-7 modelling (Panel C, P = 0.19). Panel D shows that including

State 20 produces a large and highly significant State × FSS group interaction driven entirely by the saturated 20 N condition, confirming the rationale for excluding it from the primary analysis.

**Table S2. Cohort overlap (n = 14): FSS-7 cross-session agreement details**

Subject-level FSS-7 scores from the 14 participants who completed both the IB and FMT protocols at separate testing sessions. IDs are anonymised (P1–P14, sorted by descending FSS-7 in the IB session). FSS diff is FSS-7 in the IB session minus FSS-7 in the FMT session. Category concordance indicates whether the same FSS-7 categorical assignment (LowFSS < 5; HighFSS ≥ 5) was preserved across the two sessions.

| ID | FSS-7 (IB) | Cat. (IB) | FSS-7 (FMT) | Cat. (FMT) | FSS diff | Category concordance |
|---|---|---|---|---|---|---|
| P1 | 6.71 | *High* | 5.14 | *High* | +1.57 | Concordant |
| P2 | 6.29 | *High* | 5.86 | *High* | +0.43 | Concordant |
| P3 | 6.29 | *High* | 6.00 | *High* | +0.29 | Concordant |
| P4 | 6.00 | *High* | 4.29 | Low | +1.71 | ***Discordant*** |
| P5 | 5.29 | *High* | 5.14 | *High* | +0.15 | Concordant |
| P6 | 4.86 | Low | 5.00 | *High* | -0.14 | ***Discordant*** |
| P7 | 4.00 | Low | 1.57 | Low | +2.43 | Concordant |
| P8 | 3.57 | Low | 5.71 | *High* | -2.14 | ***Discordant*** |
| P9 | 2.86 | Low | 1.57 | Low | +1.29 | Concordant |
| P10 | 2.57 | Low | 1.71 | Low | +0.86 | Concordant |
| P11 | 2.00 | Low | 3.43 | Low | -1.43 | Concordant |
| P12 | 1.57 | Low | 1.71 | Low | -0.14 | Concordant |
| P13 | 1.43 | Low | 1.00 | Low | +0.43 | Concordant |
| P14 | 1.14 | Low | 1.86 | Low | -0.72 | Concordant |

**Summary statistics.** Spearman ρ = 0.72, P = 0.004. Mean FSS-7 difference (IB − FMT) = 0.33 (SD = 1.24); 95% limits of agreement = -2.09 to 2.75. Category concordance: 11/14 (79%) of overlap participants retained the same FSS group classification across sessions; three participants switched (P4: HighFSS → LowFSS; P6 and P8: LowFSS → HighFSS), in all cases representing borderline values close to the FSS-7 ≥ 5 cut-off used throughout the analyses.

Under a pre-specified equivalence bound of half a standard deviation of the full IB cohort (+/- 0.87 FSS-7 points; rationale in Supplementary Methods, section 3), the two sessions did not differ (mean difference 0.33 points; paired t(13) = 0.99, P = 0.34), though formal equivalence was not reached at this sample size (P = 0.06).

**Table S3. Intentional binding behavioural analyses: full output and HADS sensitivity**

Type III ANOVA tables for the trial-level mixed-effects model of judgement error as a function of Judgement (Action vs Tone), Agency (Baseline vs Operant) and fatigue group, with a random intercept by participant. Panel A reports the primary categorical model (n = 23 patients); Panel B reports the same model with continuous FSS-7; Panels C and D report sensitivity analyses with HADS-Depression and HADS-Anxiety as covariates (additive and full interaction models respectively); Panel E reports the subject-level analysis of action and outcome binding indices, both categorical and continuous.

***Panel A. Primary categorical model (Judgement × Agency × FSS group)***

| Effect | Num df | Den df | *F* | *P* |
|---|---|---|---|---|
| Judgement | 1 | 3185 | 430.3 | <0.0001 |
| Agency | 1 | 3185 | 9.23 | 0.0024 |
| FSS group | 1 | 23.8 | 1.56 | 0.224 |
| Judgement × Agency | 1 | 3185 | 13.21 | <0.001 |
| Judgement × FSS group | 1 | 3185 | 14.86 | <0.001 |
| Agency × FSS group | 1 | 3185 | 0.03 | 0.872 |
| Judgement × Agency × FSS group | 1 | 3185 | 7.39 | 0.0066 |

***Panel B. Continuous FSS-7 model***

| Effect | Num df | Den df | *F* | *P* |
|---|---|---|---|---|
| Judgement | 1 | 3185 | 160.8 | <0.0001 |
| Agency | 1 | 3185 | 0.13 | 0.713 |
| FSS-7 (continuous) | 1 | 23.8 | 0.23 | 0.638 |
| Judgement × Agency | 1 | 3185 | 1.99 | 0.159 |
| Judgement × FSS-7 | 1 | 3185 | 14.26 | <0.001 |
| Agency × FSS-7 | 1 | 3185 | 0.95 | 0.331 |
| Judgement × Agency × FSS-7 | 1 | 3185 | 7.99 | 0.0047 |

***Panel C. HADS-Depression + HADS-Anxiety as additive covariates***

| Effect | Num df | Den df | *F* | *P* |
|---|---|---|---|---|
| Judgement | 1 | 3185 | 430.3 | <0.0001 |
| Agency | 1 | 3185 | 9.23 | 0.0024 |
| FSS group | 1 | 23.9 | 0.39 | 0.539 |

| Effect | Num df | Den df | *F* | *P* |
|---|---|---|---|---|
| HADS-Depression | 1 | 23.0 | 1.96 | 0.174 |
| HADS-Anxiety | 1 | 23.0 | 5.09 | 0.034 |
| Judgement × Agency | 1 | 3185 | 13.20 | <0.001 |
| Judgement × FSS group | 1 | 3185 | 14.85 | <0.001 |
| Agency × FSS group | 1 | 3185 | 0.03 | 0.872 |
| Judgement × Agency × FSS group | 1 | 3185 | 7.38 | 0.0066 |

***Panel D. HADS × FSS group interaction model***

| Effect | Num df | Den df | *F* | *P* |
|---|---|---|---|---|
| Judgement | 1 | 3185 | 430.2 | <0.0001 |
| Agency | 1 | 3185 | 9.23 | 0.0024 |
| FSS group | 1 | 23.2 | 0.01 | 0.906 |
| HADS-Depression | 1 | 23.0 | 0.23 | 0.633 |
| HADS-Anxiety | 1 | 23.0 | 2.62 | 0.119 |
| Judgement × Agency | 1 | 3185 | 13.21 | <0.001 |
| Judgement × FSS group | 1 | 3185 | 14.85 | <0.001 |
| Agency × FSS group | 1 | 3185 | 0.03 | 0.872 |
| FSS group × HADS-Depression | 1 | 23.0 | 1.28 | 0.270 |
| FSS group × HADS-Anxiety | 1 | 23.0 | 0.81 | 0.379 |
| Judgement × Agency × FSS group | 1 | 3185 | 7.38 | 0.0066 |

***Panel E. Subject-level binding indices (Action vs Outcome)***

Two-way ANOVA on the subject-level Baseline–Operant binding indices, with Binding type (Action vs Outcome) and fatigue group as factors, for the full IB cohort (n = 23). Group medians (Baseline − Operant, ms [IQR]) were: action binding, LowFSS +10.0 [34.8], HighFSS +10.3 [39.8]; outcome binding, LowFSS −18.2 [159.7], HighFSS +17.8 [70.7].

*Categorical model (Binding type × FSS group):*

| Effect | Num df | Den df | *F* | *P* |
|---|---|---|---|---|
| Binding type (Action vs Outcome) | 1 | 46.0 | 2.95 | 0.093 |
| FSS group | 1 | 46.0 | 0.00 | 0.957 |
| Binding type × FSS group | 1 | 46.0 | 1.27 | 0.265 |

*Continuous FSS-7 model (Binding type × FSS-7):*

| **Effect** | **Num df** | **Den df** | ***F*** | ***P*** |
|---|---|---|---|---|
| Binding type (Action vs Outcome) | 1 | 46.0 | 0.61 | 0.440 |
| FSS-7 (continuous) | 1 | 46.0 | 0.23 | 0.635 |
| Binding type × FSS-7 | 1 | 46.0 | 2.22 | 0.143 |

The three-way Judgement × Agency × FSS_group interaction (Panel A, $P = 0.007$) remains numerically identical with HADS-Depression and HADS-Anxiety added as additive covariates (Panel C, $P = 0.0066$) and as full FSS_group × HADS interactions (Panel D, $P = 0.0066$), confirming that the fatigue-related modulation of outcome binding is not accounted for by mood disturbance. HADS-Anxiety showed a small main effect on judgement error in the additive model (Panel C, $P = 0.034$) that did not modulate the focal three-way interaction. The subject-level binding analyses (Panel E) do not detect the fatigue-related effect, consistent with the limited power of the subject-level test ($n = 23$) and confirming the importance of the trial-level pooled analysis for detecting the effect localised to outcome binding via estimated marginal means (see main Results).

**Table S4. Readiness potential analyses: full output and HADS sensitivity**

Type III ANOVA tables for the trial-level mixed-effects models of early and late readiness potential amplitude at Cz, as a function of Agency (Baseline vs Operant) and fatigue group, with a random intercept by participant (n = 17 patients). Panels A and B concern the early RP window (−2000 to −1000 ms); Panels C and D concern the late RP window (−500 to −50 ms).

***Panel A. Early RP — Primary categorical model (Agency × FSS group)***

| Effect | Num df | Den df | *F* | *P* |
|---|---|---|---|---|
| Agency × FSS group | 1 | 2043 | 13.72 | <0.001 |

*Note. Lower-order terms (Agency, FSS group) are not estimable as independent main effects in this 2 × 2 model and are subsumed by the focal interaction. Full estimated marginal means are reported in the main Results.*

***Panel B. Early RP — HADS-Depression + HADS-Anxiety as additive covariates***

| Effect | Num df | Den df | *F* | *P* |
|---|---|---|---|---|
| Agency | 1 | 2044 | 1.93 | 0.165 |
| FSS group | 1 | 13.7 | 4.43 | 0.054 |
| HADS-Depression | 1 | 14.0 | 2.86 | 0.113 |
| HADS-Anxiety | 1 | 13.6 | 2.86 | 0.114 |
| Agency × FSS group | 1 | 2045 | 13.62 | <0.001 |

***Panel C. Late RP — Primary categorical model (Agency × FSS group)***

| Effect | Num df | Den df | *F* | *P* |
|---|---|---|---|---|
| Agency × FSS group | 1 | 2039 | 8.26 | 0.0041 |

***Panel D. Late RP — HADS-Depression + HADS-Anxiety as additive covariates***

| Effect | Num df | Den df | *F* | *P* |
|---|---|---|---|---|
| Agency | 1 | 2038 | 8.23 | 0.0042 |
| FSS group | 1 | 15.6 | 0.94 | 0.347 |
| HADS-Depression | 1 | 15.8 | 2.01 | 0.176 |
| HADS-Anxiety | 1 | 15.6 | 1.37 | 0.259 |
| Agency × FSS group | 1 | 2039 | 8.23 | 0.0042 |

***Panel E. Continuous FSS-7 models (Agency × FSS-7), early and late RP***

| Window | Effect | Num df | Den df | *F* | *P* |
|---|---|---|---|---|---|
| Early RP | Agency | 1 | 2039 | 16.10 | <0.0001 |

| Window | Effect | Num df | Den df | *F* | *P* |
|---|---|---|---|---|---|
| Early RP | FSS -7 (continous) | 1 | 15 | 0.28 | 0.605 |
| Early RP | Agency × FSS-7 | 1 | 2039 | 9.64 | 0.002 |
| Late RP | Agency | 1 | 2037 | 14.22 | 0.0002 |
| Late RP | FSS -7 (continous) | 1 | 16 | 0.02 | 0.888 |
| Late RP | Agency × FSS-7 | 1 | 2037 | 5.64 | 0.018 |

The Agency × FSS group interaction is essentially unchanged under HADS adjustment for both the early RP window ($P = 0.0002$ without adjustment vs $P = 0.0002$ with HADS adjustment) and the late RP window ($P = 0.0041$ without adjustment vs $P = 0.0042$ with HADS adjustment), confirming that the fatigue-related disruption of agency-related readiness-potential modulation is not accounted for by mood disturbance. Neither HADS-Depression nor HADS-Anxiety showed a significant main effect on early or late RP amplitude (all $P > 0.11$). The interaction is also significant under continuous FSS-7 modelling at both windows (early: $P = 0.002$; late: $P = 0.018$; Panel E), confirming that the effect is not an artefact of the categorical FSS split.

**Table S5. Robustness, equivalence and power analyses**

Additional analyses conducted to strengthen inference given the modest sample sizes. Part (a) reports robustness of the early readiness-potential interaction (Supplementary Methods, section 7); part (b) the direct between-subject test of predictive-postdictive uncoupling (section 6); part (c) the minimum detectable effect and equivalence testing for the force-matching interaction (section 8).

| Analysis | Estimate / statistic | P-value or interval |
|---|---|---|
| **Panel A. Early readiness-potential interaction robustness (n = 17)** | | |
| Permutation test (5000 permutations, subject-level shuffle) | — | P = 0.0004 |
| Leave-one-out refitting (17 fits) | F range 9.12–14.14 | significant in 17/17 (P < 0.05) |
| Bayesian interaction coefficient | +2.12 μV | 95% CrI 0.98 to 3.25; P(direction) > 0.99 |
| Standardised Baseline−Operant contrast, LowFSS | d = 0.25 | 95% CI 0.14 to 0.35 |
| Standardised Baseline−Operant contrast, HighFSS | d = −0.11 | 95% CI −0.27 to 0.05 |
| **Panel B. Predictive–postdictive uncoupling (n = 17)** | | |
| Partial Spearman (controlling FSS-7) | ρ = −0.16 | P = 0.56 |
| Raw Spearman (group-confounded) | ρ = −0.19 | P = 0.47 |
| **Panel C. Sensory-attenuation minimum detectable effect and equivalence** | | |
| Minimum detectable interaction slope, 80% power | 0.28 N per force level | — |
| Minimum detectable interaction slope, 90% power | 0.32 N per force level | — |
| Mean SA, equivalence (±0.5 SD) | t (12.62) = −0.32 | P = 0.75; equivalence P = 0.18 |
| Gradient, equivalence (±0.5 SD) | t (25.37) = 1.47 | P = 0.15; equivalence P = 0.47 |
| Intercept, equivalence (±0.5 SD) | t (9.92) = −0.76 | P = 0.46; equivalence P = 0.35 |

### Supplementary Figures

#### Figure S1. Judgement error distributions per condition

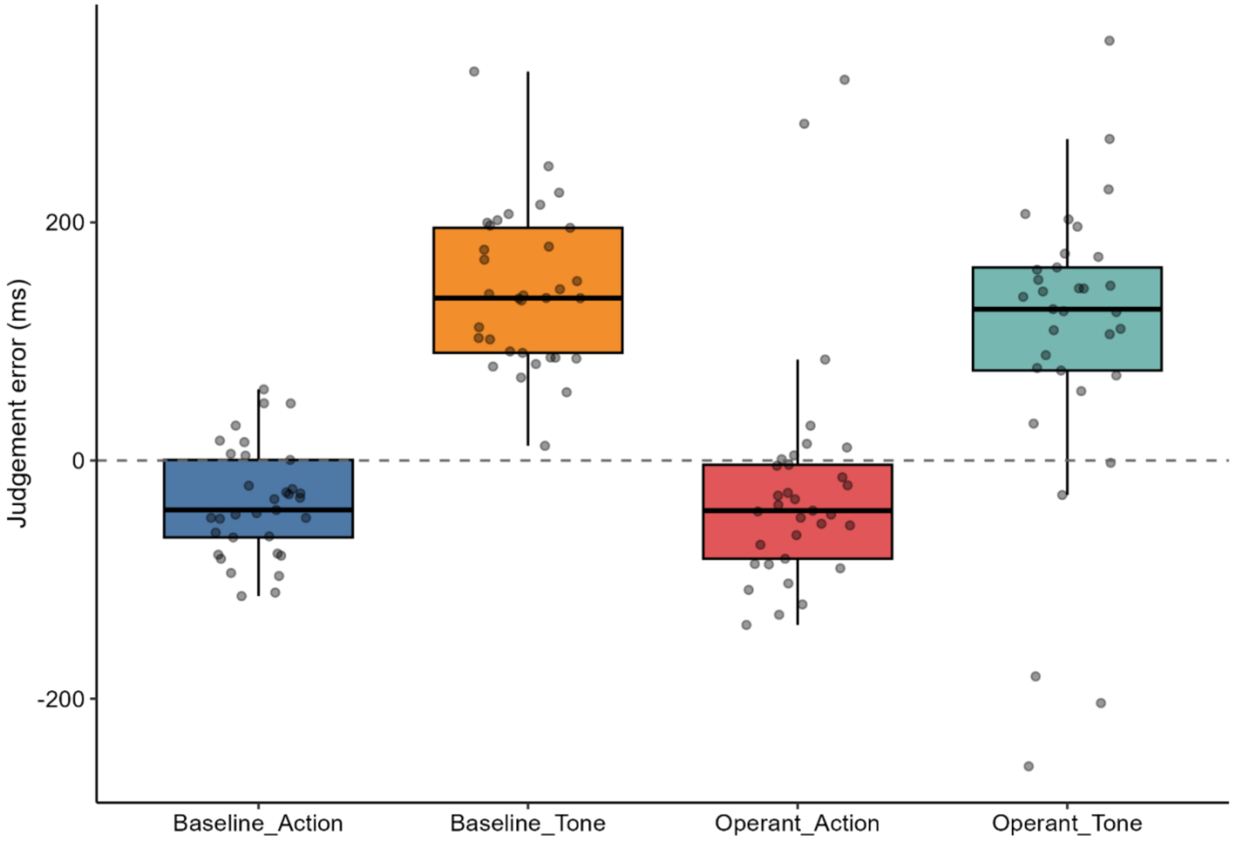


**Figure S1.** Subject-level median judgement error (ms) for each of the four conditions (Baseline Action, Operant Action, Baseline Tone and Operant Tone) in the full Intentional Binding cohort (n = 23). Each point represents one participant; boxplots show the median and interquartile range, with whiskers extending to 1.5 × the interquartile range. Judgement error is the reported minus the actual event time (ms); positive values indicate the event was perceived as occurring later than it actually did, negative values earlier. The dashed line marks veridical timing (zero error). This figure displays the raw distributions underlying the trial-level mixed-effects model reported in the main Results (Figure 4).

**Figure S2. Subject-level binding distributions stratified by fatigue group**

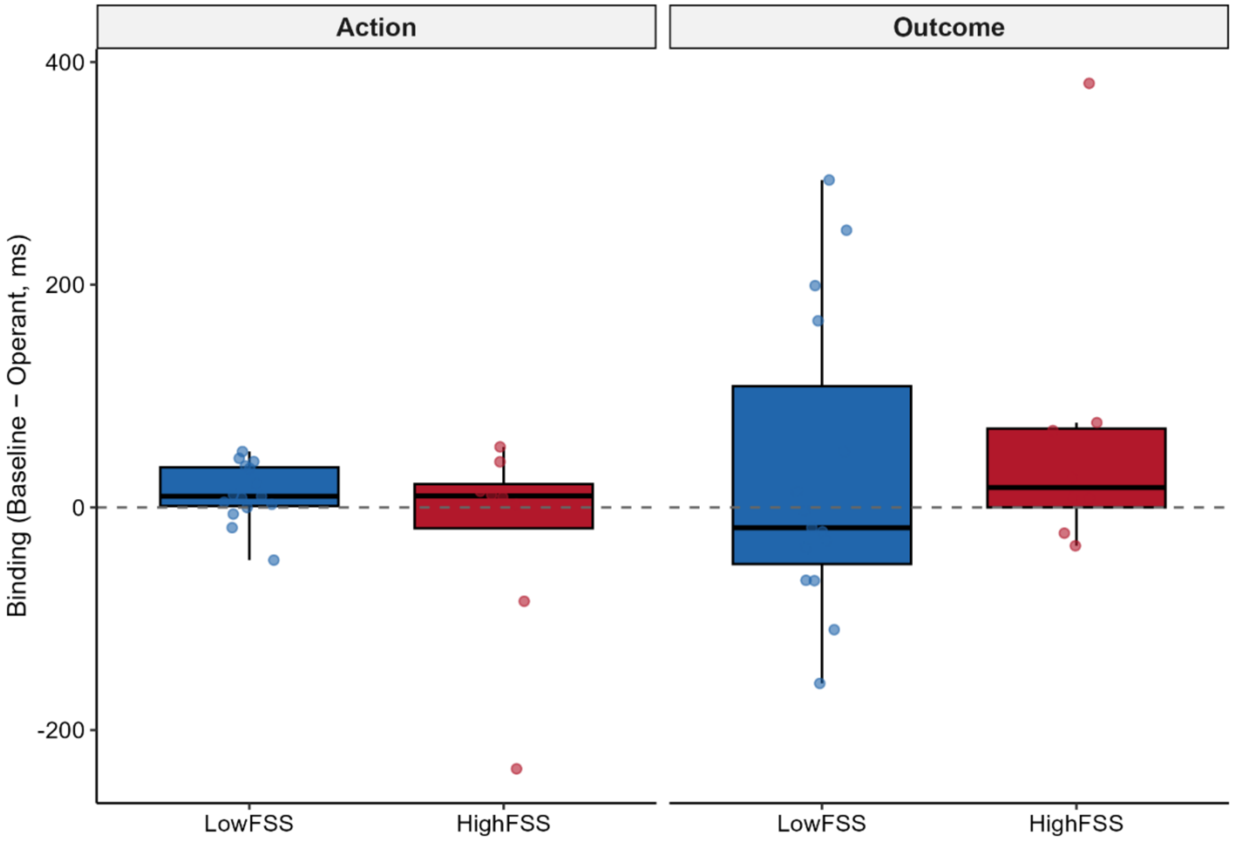


**Figure S2**. Subject-level Baseline–Operant binding indices for the Action and Outcome components, shown separately for low-fatigue (LowFSS, n = 15) and high-fatigue (HighFSS, n = 8) patients. Each point represents one participant; boxplots show the median and interquartile range. Action binding is computed as the Baseline–Operant shift in the Action judgement and outcome binding as the corresponding shift in the Tone judgement, such that positive outcome-binding values indicate the canonical compression of perceived outcome timing toward the action. These subject-level distributions complement the trial-level estimated marginal means reported in the main Results (Figure 4). The corresponding subject-level analysis is reported in Supplementary Table S3, Panel E.

**Figure S3. Force matching task: matched vs applied force scatter per group**

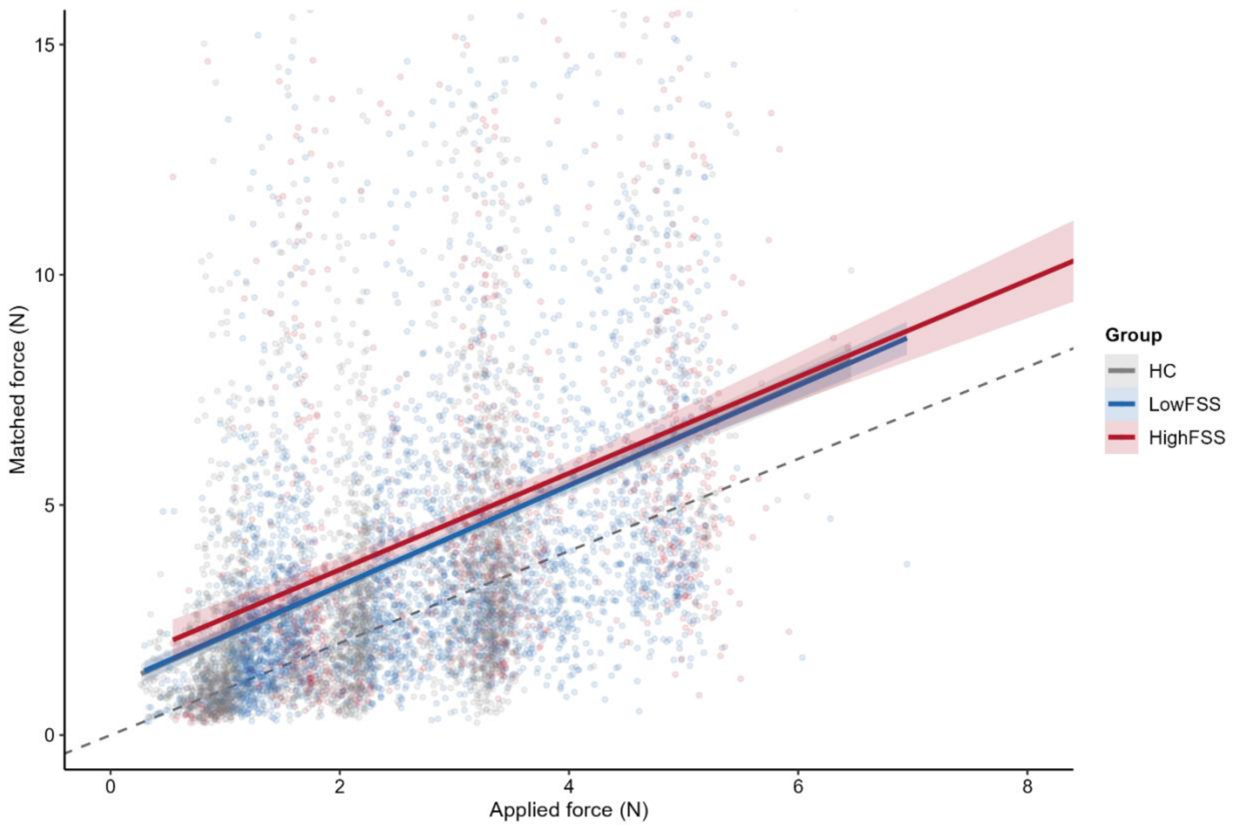


**Figure S3.** Single-trial matched force (y-axis) plotted against applied target force (x-axis), pooled across target levels, for healthy controls (HC, grey), low-fatigue patients (LowFSS, blue) and high-fatigue patients (HighFSS, red). Faint points are individual trials. Solid lines are ordinary least-squares fits per group with 95% confidence bands (shaded). The dashed line is the line of identity (matched = applied): fitted lines lying above it denote overcompensation of force (matched > applied), the behavioural index of sensory attenuation in this paradigm (Shergill et al., 2003). All three group fits lie above the identity line across the sampled force range. The LowFSS and HighFSS fits overlap closely throughout the densely sampled range, their divergence at the upper end reflecting the few high-force HighFSS trials. This figure is a descriptive complement to the formal between-group comparison of matching behaviour, reported in the main-text FMT analysis, which showed no significant group differences in mean attenuation, intercept or gradient.

N = HC 24; LowFSS 30; HighFSS 10, after primary quality control (Supplementary Methods 2).

**References (Supplementary Materials only)**


1. Shergill SS, Bays PM, Frith CD, Wolpert DM. Two eyes for an eye: the neuroscience of force escalation. *Science*. 2003;301(5630):187. doi:10.1126/science.1085327

2. Norman GR, Sloan JA, Wyrwich KW. Interpretation of changes in health-related quality of life: the remarkable universality of half a standard deviation. *Med Care*. 2003;41(5):582-592. doi:10.1097/01.MLR.0000062554.74615.4C

3. Bates D, Mächler M, Bolker B, Walker S. Fitting Linear Mixed-Effects Models Using lme4. *Journal of Statistical Software*. 2015;67:1-48. doi:10.18637/jss.v067.i01

4. Kuznetsova A, Brockhoff PB, Christensen RHB. lmerTest Package: Tests in Linear Mixed Effects Models. *Journal of Statistical Software*. 2017;82:1-26. doi:10.18637/jss.v082.i13

5. Lenth R, Piaskowski J. emmeans: Estimated Marginal Means, aka Least-Squares Means. *mmeans: Estimated Marginal Means, aka Least-Squares Means R package version 203,*. Published online 2026. Accessed July 8, 2026. https://rvlenth.r-universe.dev/emmeans

6. Jo HG, Wittmann M, Hinterberger T, Schmidt S. The readiness potential reflects intentional binding. *Front Hum Neurosci*. 2014;8:421. doi:10.3389/fnhum.2014.00421

7. Kim S. ppcor: An R Package for a Fast Calculation to Semi-partial Correlation Coefficients. *Commun Stat Appl Methods*. 2015;22(6):665-674. doi:10.5351/CSAM.2015.22.6.665

8. Bürkner PC. brms: An R Package for Bayesian Multilevel Models Using Stan. *Journal of Statistical Software*. 2017;80:1-28. doi:10.18637/jss.v080.i01

9. Lakens D. Equivalence Tests: A Practical Primer for t Tests, Correlations, and Meta-Analyses. *Social Psychological and Personality Science*. 2017;8(4):355-362. doi:10.1177/1948550617697177

10. Caldwell A. Exploring Equivalence Testing with the Updated TOSTER R Package. Published online November 17, 2022. Accessed July 8, 2026. https://osf.io/ty8de_v1
